# Temporal Structure of Music Improves the Cortical Encoding of Speech

**DOI:** 10.1101/2024.06.14.598982

**Authors:** Laura Fernández-Merino, Mikel Lizarazu, Nicola Molinaro, Marina Kalashnikova

**Affiliations:** Basque Center on Cognition, Brain and Language, San Sebastian, Spain; Ikerbasque. Basque Foundation for Science, Bilbao, Spain; University of the Basque Country (Universidad del País Vasco/Euskal Herriko Unibertsitatea), San Sebastian, Spain

**Keywords:** Cortical tracking, rhythm, musical exposure, oscillations

## Abstract

Long and short-term musical training has been proposed to improve efficiency of cortical tracking of speech, the mechanism through which brain oscillations synchronize to the acoustic temporal structure of external stimuli. Here, we study how different rhythm structures of the musical signal can guide the temporal dynamics of auditory oscillations phase-aligned to the speech envelope. For this purpose, we investigated the effects of prior exposure to rhythmically structured musical sequences on cortical tracking of speech in Basque-Spanish bilingual adults. We conducted two EEG experiments where participants were presented with sentences in Basque and Spanish preceded by musical sequences that differed in their beat structure. The beat structure of the musical sequences was created to 1) reflect and match the syllabic structure of the sentences, 2) reflect a regular rhythm but not match the syllabic structure of the sentences, and 3) follow an irregular rhythm. First, we showed that the regularity found in the rhythmic structure of music acts as a temporal guide for brain oscillations. Second, our findings suggest that not only the regularity in music is crucial but so is adjusting this regularity to optimally reflect the rhythmic characteristics of the language. Third, despite finding some differences across frequencies for each language, we still found a strong effect of rhythm regularity on cortical tracking of speech. We showed that rhythm, inherent in musical signals, guides the adaptation of brain oscillations, by adapting the temporal dynamics of the oscillatory activity to the rhythmic scaffolding of the musical signal.

## Introduction

There is extensive evidence that experienced musicians and even individuals with more limited music training experience cognitive advantages when processing speech (see Milovanov & Tervaniemi, 2011 for review; also, Patel and Iversen, 2007). However, little is known about the exact mechanisms underlying these benefits. Recently, attention has been paid to the role of musical stimulation on auditory cortical tracking, which is the mechanism through which the brain oscillations synchronize to the acoustic properties of external stimuli. From this perspective, the precise aspects of musical stimulation that may lead to these music-to-speech cortical tracking effects are yet unknown. Here, we propose that rhythmic auditory stimulation, as found in the musical signal, acts as a temporal guide that helps to maintain the temporal dynamics of the synchronization to the speech signal over time.

Music and language communication are human cultural universals that are present from very early in our lives (Peretz, 2006). There is a growing body of literature that has tried to establish links between music and language processing in the brain, especially music-to-speech processing benefits (see Besson et al., 2017 for a review). Research involving professional musicians has shown that music experience enhances understanding speech in noise and sensitivity to prosodic cues (Zendel et al., 2019). More specifically, music training improves auditory processing in noisy conditions by strengthening the sensorimotor coupling between premotor and auditory regions (Du & Zatorre, 2017; Herholz, & Zatorre, 2012). This, in turn, is related to finer analysis of the spectral properties of the auditory signal, possibly due to greater neural temporal precision (Tierney, & Kraus, 2013; Wong et al., 2007), and has positive effects on audiovisual speech integration (Musacchia et al., 2007).

There is also evidence showing that these processing benefits are observed in non-musical experts after short-term music training. Moreno and Besson (2006) found that children attending eight weeks of music training showed a decrease in the amplitude of late positive components to strong incongruities in pitch compared to children attending painting training. In a longitudinal study following six months of either musical or painting training, Moreno et al. (2009) showed that children in the musical training group presented improved reading and pitch discrimination abilities in speech. In a study with adolescents, Tierney et al. (2013) demonstrated that two years of music instruction improved the neural representation of speech in noisy environments. Along similar lines, three years of music instruction proved to enhance subcortical sound processing and to accelerate the maturation of cortical auditory responses (Tierney et al., 2015). Furthermore, other studies have shown that these benefits extend to more general language abilities such as vocabulary skills (Piro & Ortiz, 2009), phonological awareness (Dege & Schwarzer, 2011), and auditory processing (Habibi et al., 2016), as shown in children and adults, as well as young infants (Gerry et al., 2012; Zhao & Kuhl, 2016).

Based on this growing of evidence, some theories have outlined shared underlying processes for music and speech processing that may explain why there is an improvement in speech processing from music exposure. The Overlap, Precision, Emotion, Repetition, and Attention (OPERA) hypothesis (Patel, 2011; 2014), proposes that music training enhances speech processing when music places higher demands than speech on shared cognitive processes, and engages this process in the context of emotion, repetition, and attention. The Precise Auditory Timing Hypothesis (PATH; Tierney & Kraus, 2014) explains enhanced linguistic abilities in musicians by cross-domain auditory improvement; in other words, changes in neural processing in one domain are driven by experience or training in another. While both these frameworks suggest potential transfer effects across domains, this transfer seems to be driven by experience or long-term music training. However, little is known about why and how non-musicians would benefit from short-term musical exposure.

It has also been proposed that there is a shared nature of structural processing in language and music - including rhythm processing (Fiveash et al., 2021). Language and music are hierarchically organized, and as such, one of the main foci of the last few years has been to establish the neurobiological underpinnings of how the human brain processes these hierarchies. One of the perspectives that has been adopted to study this hypothesis posits that neural activity synchronizes to the rhythmical patterns of external inputs, such as speech and musical signals (Obleser & Kayser, 2019). This synchronization mechanism, referred to as *cortical tracking* hereafter, is reflected in the phase and amplitude of the neural oscillations that synchronize with the stimulus (see Doelling & Assaneo, 2021 for review). Cortical tracking of auditory stimuli plays a role in processing the linguistic structure of speech, via aligning specific neural oscillators to specific linguistic units: phonological rate (gamma, >30 Hz) (Hyafil et al., 2015), syllabic rate (theta, 4-8 Hz) (Peelle et al., 2013), and lexical and phrasal rate (delta, 1-3 Hz) (Bourguignon et al., 2013; Kong et al., 2015). The tracking of the incoming signal allows brain oscillations to synchronize with external regularities and predictable cues (e.g., beat in music or stress in speech), and to structure the auditory input. It also helps focus attention on important elements of the auditory stimulus and its presentation over time (Ghitza, 2011; Giraud & Poeppel, 2012; Peelle & Davis, 2012), supporting the segmentation and identification of linguistic units in speech (Meyer, 2018). Accurate tracking of acoustic temporal regularities correlates with speech intelligibility (Ahissar et al., 2001; Luo & Poeppel, 2007), and has been suggested to be responsible for accurate syllabic parsing (Doelling et al., 2014). In contrast, diminished cortical tracking has been found in challenging listening conditions (Peelle et al., 2013). While it is well established that cortical tracking underlies the perception of rhythmic regularities in speech and music, whether and how this mechanism modulates the music-to-speech processing benefits is less clear.

Music and speech share a structure and hierarchy with a common underlying denominator – rhythm (Ding et al., 2017; Varnet et al., 2017). A growing body of literature that recognizes the importance of rhythm for language acquisition, speech, and music processing (Goswami, 2022; Ladányi et al., 2020). From the perspective of cortical tracking, for instance, delta brain oscillations phase-align with the beats of the rhythm of speech and music (Lakatos et al., 2013; 2016; Poeppel & Assaneo, 2020; Schroeder & Lakatos, 2009). In addition, rhythmic priming studies suggest that the temporal regularity found in music or speech rhythms would enable prediction of when upcoming beats or syllables will occur, and that results in enhanced perception at the times of beat and syllable onsets. In such studies, participants have been shown to benefit from congruent rhythmic primes, when the syllabic structure of the subsequent pseudo-words matched the rhythmic prime metrical structure in phonological (Cason & Schön, 2012), prosodic (Ríos-López et al., 2017), and syntactic perception (Przybylski et al., 2013). These studies suggest that rhythmic regularity, through rhythmic priming, can prime subsequent processing by enhancing the phase of oscillations, thereby enhancing cortical tracking.

Starting from the assumption that cortical tracking is the mechanism through which language processing is enhanced in individuals when exposed to music, here we propose that the rhythmic scaffolding of the music signal boosts the adaptation of the oscillatory activity to the temporal dynamics of speech. Critically, we propose that music-to-speech benefits arise from the similarities between the temporal structure of music and speech. Specifically, from the rich and complex structure in music reflecting the prosodic and syllabic rhythms of speech. In line with this prediction, one previous study has found enhanced cortical tracking of speech in non-musical experts following exposure to speech-to-musical rhythms. Falk et al (2017) showed that exposure to regular musical cues that reflect the temporal and prosodic structure of speech enhances inter-trial phase consistency during speech listening. In this electroencephalography (EEG) study, participants’ neural responses were recorded while they listened to speech sequences (participants were French speakers listening to French speech) preceded by musical sequences that differed in rhythm regularity. Speech stimuli were marked by pitch and duration variations, modulating syllabic rate at 5 Hz. Musical sequences either had regular rhythm, i.e., the metrical structure was identical to the temporal structure of speech, or irregular rhythm, i.e., the metrical structure did not follow any regular rhythm and was different from the temporal structure of speech. Regular musical sequences’ modulations were prominent at the meter level, 1.65 Hz, and beat level, 5 Hz, reflecting the prosodic and syllabic rhythms of the speech sequences. Inter-trial phase consistency analyses showed larger phase consistency to speech in the regular condition compared to the irregular condition, i.e., when the temporal structure of speech and the preceding music sequences coincided. As mentioned, the musical sequences were reflecting the syllabic and prosodic rhythms in the delta (∼2Hz) and theta (∼5Hz) frequency bands. Participants showed larger phase consistency in the regular condition in the theta frequency band compared to the delta frequency band. This is not an unexpected effect given the rhythmical properties of French – a syllable-timed language, which modulations are highlighted in the theta frequency band.

Falk et al.’s (2017) study is important as it showed the music-to-speech cortical tracking effect can arise within a single experimental session. However, the rhythmical patterns of the music were exactly timed with the speech stimuli, which may have been key for the music-to-speech phase consistency effect. Therefore, it is possible that the improvement effect stemmed from the regular music rhythms preparing the brain to synchronize with a similar temporal structure in the following speech sequences. These findings suggest that to best synchronize to the acoustic stimuli, brain oscillations need optimal matching conditions (i.e., music and speech with the same temporal structure). However, it is also possible that the oscillations adapt to a wide range of rhythmic stimuli, and that the specific input properties required to improve this mechanism are not rigidly defined. In this study, we question whether a sequence of beats that mimics the prosodic structure of speech would be sufficient to improve cortical tracking of speech. That is, whether cortical tracking of speech can be improved when speech is preceded by a musical sequence that has a regular rhythm, but which metrical structure is not identical to the syllabic speech temporal structure. If so, the music-to-speech cortical tracking effect would be the consequence of the stimulation of the prosodic rhythms of speech, as reflected in the music.

The overall aim of the present study was to understand the precise aspects of music stimulation that lead to music-to-speech benefits via enhancing cortical tracking. To this end, we used electroencephalography (EEG) to measure cortical tracking of speech sequences preceded by musical sequences. We manipulated the rhythmical properties of the musical stimuli according to the syllabic and prosodic rates of the speech sequences, creating three conditions: Matching Regular sequences, which matched the syllabic, and therefore, temporal structure of the sentences; Mis-matching regular sequences, which had a regular rhythm but did not match the syllabic structure of the sentences; and Irregular sequences, which followed an irregular rhythm. With this study, we also aimed to clarify the relevance of the music-matching rhythms structure for the efficacy of musical stimulation on cortical tracking. The Mis-matching regular condition allows to test the role of the matching between the syllabic structure and the beat structure of the musical stimuli on speech processing. Again, musical cues in this condition follow a regular rhythmical pattern, but their beat structure is not matched to the syllabic structure of the subsequent speech stimuli. If the acoustic input drives the neural activity in the auditory cortex, we predict that neurons’ phase synchronization to rhythmical regularities would drive the music-to-speech benefit, indicating that rhythm regularity is sufficient to enhance cortical tracking of speech. Thus, we would expect higher cortical tracking of speech when preceded by both Matching and Mis-matching regular sequences than when preceded by Irregular sequences, regardless of the rhythmical differences between these conditions. If, however, the music-to-speech cortical tracking effect is specifically related to the similarities of the temporal structure between speech and music, these similarities might be crucial. In this case, we would expect higher cortical tracking to speech when preceded by Matching regular sequences compared with when speech is preceded by both Mis-matching regular and Irregular musical sequences.

In addition, we capitalized on the balanced bilingual status (Basque-Spanish) of our participants to incorporate a secondary aim, namely, to identify the role of language-specific rhythms on the music-to-speech cortical tracking effect. While Basque and Spanish are both syllable-timed languages, Basque also exhibits the characteristics of stress-timed languages due to its syllabic and syntactic structure (Molnar et al., 2016; Molnar et al., 2014). By studying how the cortical tracking effect manifests in both these languages we can investigate the generalizability of this effect across different language types. This also enables us to assess the potential impact of unique rhythmic structures on cortical tracking of speech, and its manipulation through music-to-speech rhythms. To this end, the rhythmical properties of the musical stimuli were manipulated according to the syllabic and prosodic rates of Basque and Spanish. We expected to find a music-to-speech cortical tracking effect across participants in both the Spanish and Basque tasks.

That is, participants will show enhanced cortical tracking of Spanish and Basque speech sequences after being exposed to musical sequences reflecting the specific languages’ temporal structure. However, the first experiment reported below revealed that the speech sequences designed for Basque did not accurately reflect the prosodic and syllabic rhythms of the language. Therefore, we also report a second experiment in which we fine-tuned the speech and musical stimuli in Basque and collected new data from a subset of participants.

In view of the robust effect of musical training and expertise on cortical tracking of speech discussed above, we also collected information about participants’ musical background to explore the relationship between prolonged musical training and the music-to-speech cortical tracking effect in our sample. We expect participants with frequent exposure to music to show an improvement of cortical tracking to speech in the Matching and Mis-matching regular conditions.

## Experiment 1

### Method

#### Participants

A total of 33 Basque-Spanish bilinguals (11 male, *M* age: 25 years, *SD* = 3.31) recruited through the Basque Center on Cognition, Brain, and Language (BCBL) participant database took part in this study. Language proficiency was assessed with the Basque, English, and Spanish Test (BEST) (De Bruin et al., 2017), which measures fluency, lexical resources, grammatical constructions, and pronunciation (a Likert scale with scores ranging from 1 to 5). All participants were highly proficient in Basque and Spanish (Maximum score: 65; Spanish *M:* 64, *SD*: 1.01; Basque *M*: 58.09, *SD* = 5.95). Participants’ musical background was assessed using the Gold MSI self-report questionnaire, which measures different aspects of musical sophistication, such as active musical engagement, perceptual abilities, musical training, singing abilities, emotional engagement with music, providing a general musical sophistication factor score (Müllensiefen et al., 2014). All participants self-reported to be actively engaged with music: 7 participants had more than 5 years of formal musical education, but no participant reported to be a professional musician (general musical sophistication factor, maximum score = 100; *M* = 66.45; *SD* = 11.37).

The Ethics and Scientific Committee of the BCBL approved the study protocol (approval number 220321SM), which was developed following the declaration of Helsinki. All participants gave their written informed consent prior to the study and received monetary compensation.

#### Stimuli

Speech stimuli consisted of 64 12-syllable sentences in both Spanish and Basque (128 in total) recorded by a female Basque-Spanish bilingual speaker. We used stimuli recorded in infant-directed speech (IDS) for an identical study with infants because this register provides the ideal context to stress and highlight the temporal modulations of speech (Leong et al., 2017) and ensures natural and highly rhythmical stimuli that highlighted the syllable information.

This experimental design required the creation of three types of musical stimuli for the Matching Regular, Mis-matching regular, and Irregular conditions. The musical stimuli for the Matching regular and Mis-matching regular conditions were created based on the temporal structure of the sentences. The Matching Regular musical sequences reflected the temporal structure and pitch contour of linguistic stimuli. For this purpose, we extracted the rhythmic structure of the sentences to create the beat structure. This structure was introduced in Musescore 3 (https://musescore.org/es), an open-source software where, using a grand piano timber, we created the melodic contour from the pitch contour of the sentences. The Mis-matching regular musical sequences preserved the pitch contour of the linguistic stimuli but followed a temporal structure with beats that did not match the syllabic structure of the speech stimuli. To create these sequences, we took the Matching Regular sequences and altered the duration of the beats, preserving a regular meter. Finally, the Irregular musical sequences were created by altering the pitch and note duration of the Matching Regular sequences, thus also altering their regular meter. The melodic sequences that comprised a condition formed a musical structure of A A A A’ (A’ was a slight variation of the melodic contour and temporal structure of A), following the common musical pattern of Basque and Spanish infant-directed songs.

All speech and melodic sequences had a duration of 2500 msec. To ensure highly rhythmical stimuli, each speech sequence was recorded cued by a metronome at regular rate of 120 bpms, reflecting the rate of stressed syllables, obtaining a regular meter with an inter-onset interval of 500 msec between accented syllables. The musical sequences were created based on the temporal patterns of the sentences. There was a constant 500 msec inter-onset interval between accented beats for the Matching Regular and Mis-matching Regular musical sequences, and no constant inter-onset interval for the Irregular sequences.

Given Spanish and Basque have different rhythmical properties (Molnar et al., 2016, Molnar et al., 2014), spectral analyses of the melodic and speech sequences revealed different frequencies for each language, with peaks at 2 Hz and 4 Hz in Spanish and peaks at 1.6 Hz and 4 Hz in Basque, as shown in Figure 1. These peaks were expected because they correspond to the prosodic (reflected in the delta frequency band, ∼2Hz) and syllabic (reflected in the theta frequency band, ∼4Hz) rhythms of Spanish and Basque. Thus, the difference in Matching Regular and Mis-matching Regular musical sequences remained in the matching between beats and syllables. No specific frequency was reflected in the Irregular musical sequences.

**Figure 1.**
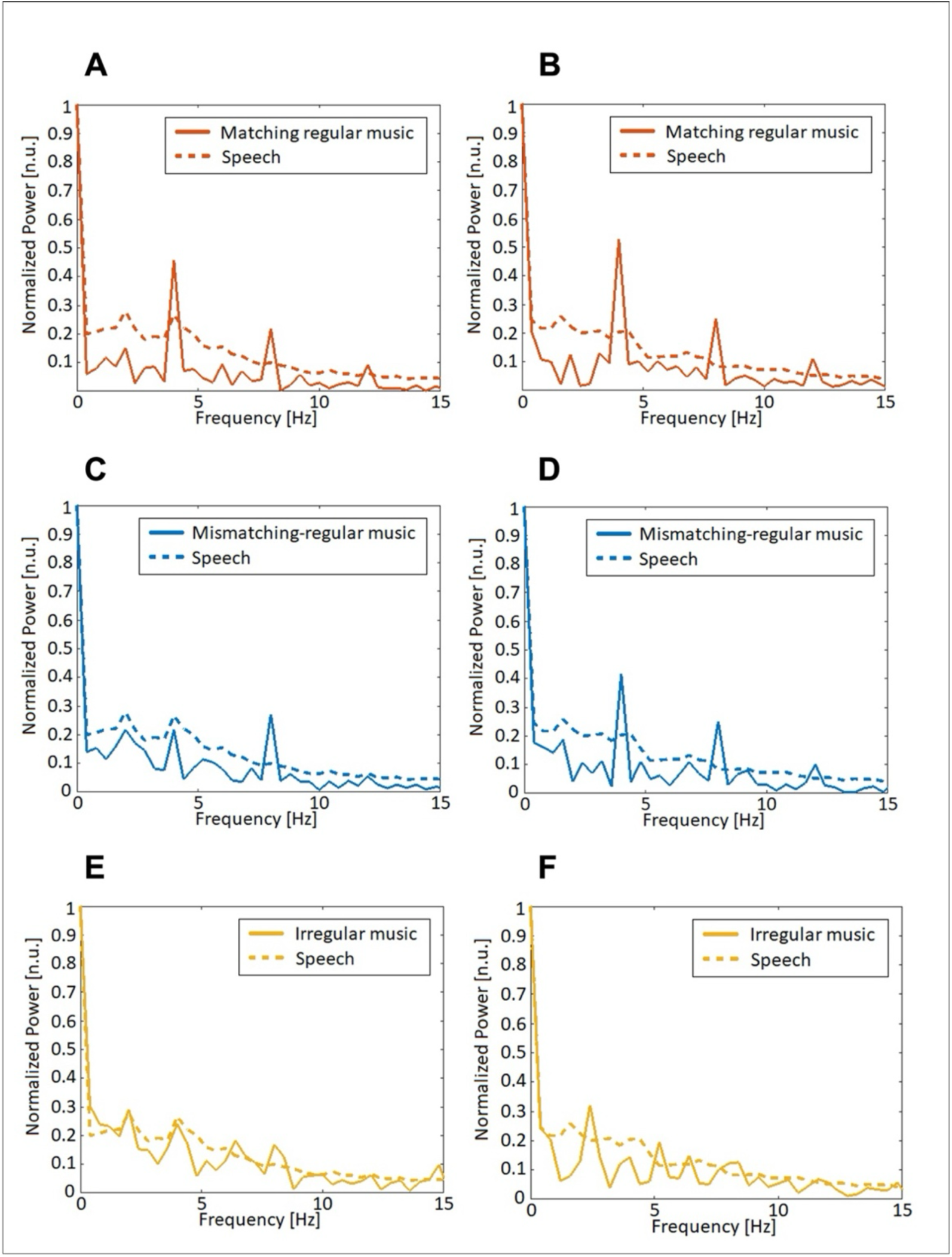
Acoustic properties of Spanish and Basque musical and speech sequences in Experiment 1. Here we show the spectral information of speech and musical sequences in normalized power (scaled power of signal). Spectra on the left (A C E) correspond to Spanish stimuli, showing peaks at 2 Hz and 4 Hz for the Matching and Mis-matching Regular condition. Spectra on the right (B D F) correspond to Basque stimuli, showing peaks at 1.6 Hz and 4 Hz.

#### Procedure

Participants completed all three conditions in Spanish and Basque in a single experimental session: speech preceded by Matching Regular musical sequences speech preceded by Mis-matching Regular cues, and speech preceded by Irregular musical cues. Stimulus presentation was structured in 16 blocks of four trials. Block structure is shown in Figure 2. Only one type of cue, Matching Regular, Mis-matching Regular, and Irregular, and one language were used within a single experimental block. There was a total of 96 blocks, with a total experiment duration of 32 minutes. There were 16 blocks per language (Basque or Spanish) and per rhythm condition (Matching, Mis-matching, and Irregular), each condition had a duration of ∼5’30”. All linguistic stimuli were preceded by Matching Regular musical sequences Mis-matching regular musical sequences, and Irregular musical sequences. A single block consisted of four trials and had a total duration of 20 sec. Each trial comprised one music sequence immediately followed by a simple sentence and had a duration of 5000 msec. The stimulus onset asynchrony (SOA) between the melodic sequence and the sentence was always 500 msec.

**Figure 2.**
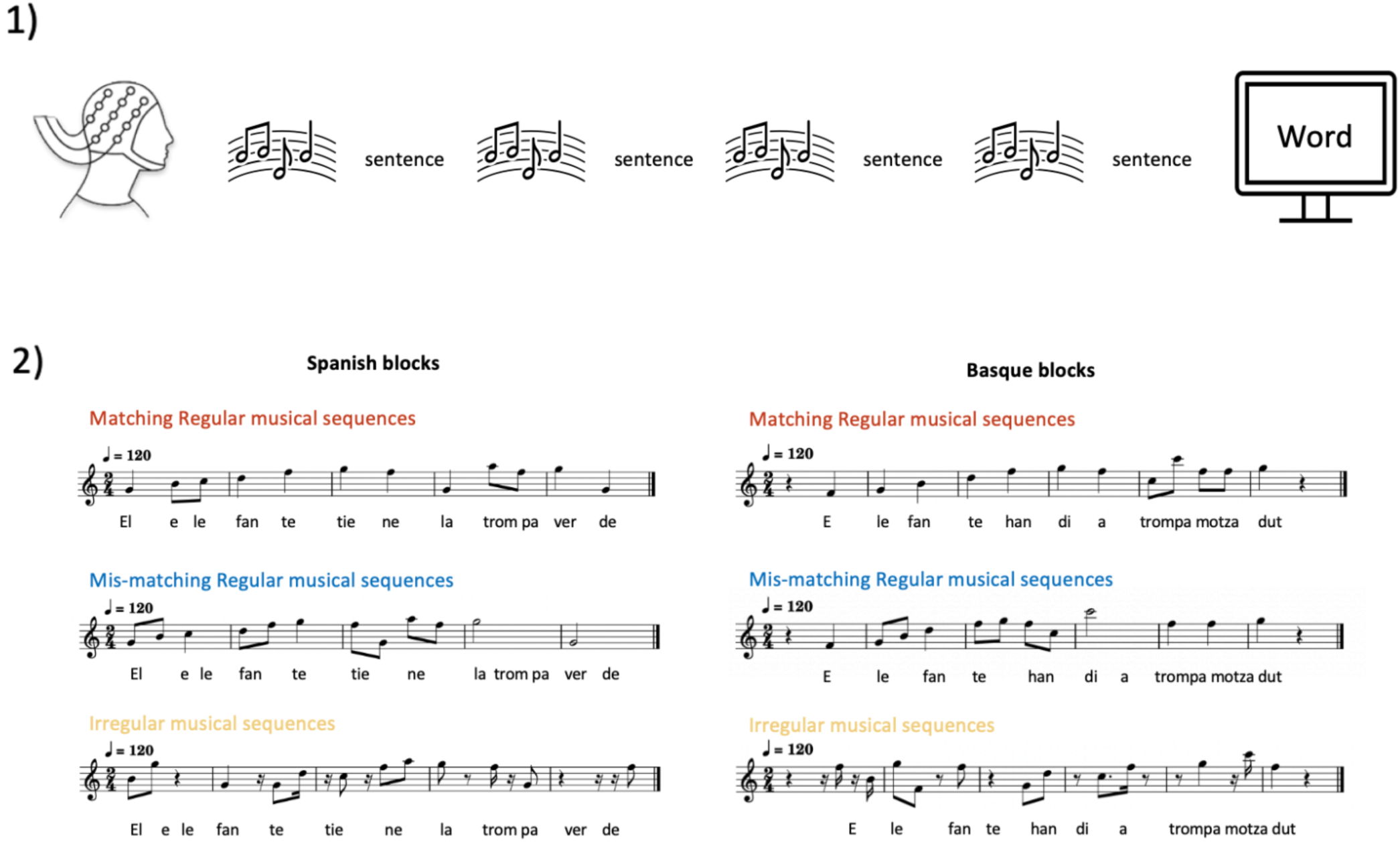
Procedure. 1) Graphical example of an experimental block. The block comprises four trials. Each trial comprises one melodic sequence followed by a sentence. Only one type of cue (Matching, Mismatching or Irregular was used within a block). After each block, a word appeared on the screen. 2) Example of the musical sequences for each condition. Below each pentagram, an example of the sentence that would follow the musical sequences (English translation: the elephant’s trunk is green). Note that only in the Matching Regular musical sequences, the beat structure of the musical sequences and the syllabic structure of the sentences are matched.

To ensure participants’ attention during data acquisition, after each block, a word appeared on the screen and participants had to indicate, by pressing a key, whether they had or had not heard that word in the sentences in that block. Half of the words appeared in the sentences and half of the words did not. The next block started immediately after a key was pressed.

The order of condition and language was counterbalanced across participants. Stimuli were presented in PsychoPy (v.1.80.04) via two speakers positioned about one meter in front of the participants, at 65 dB.

For all participants we also recorded resting state EEG data, as participants looked at a fixation point during five minutes at the end of the experimental session.

#### EEG data acquisition and preprocessing

EEG data were acquired using a BrainAmp amplifier and BrainVision Recorder software (Brain Products, Germany). EEG was recorded using 32 electrodes that were positioned according to the international 10–20 system (Jasper, 1958). Scalp-electrode impedance was kept below 5 kΩ for scalp electrodes and under 10 kΩ for reference and EOG electrodes to ensure high-quality EEG recordings. EEG data was sampled at a rate of 1000 Hz and band-pass filtered online from 0.1 to 1000 Hz. The recording was referenced to electrode FCz online. Electrode AFz was used as the ground. Additionally, two electrodes at the outer canthi of the eyes recorded horizontal eye movements, while electrodes above and below the right eye recorded vertical eye movements.

All the following processing and analysis steps were implemented in Matlab (The MathWorks, Inc., Natick, US) using FieldTrip (Oostenveld et al., 2011) and available custom code (Fernández-Merino & Lizarazu, 2023). Data were segmented from 0. s to 2.5 s relative to stimulus onset, which gave a total of 768 trials. These trials were re-referenced off-line to the left mastoid and low pass filtered below 30 Hz, since we did not expect any coherence effect above this threshold (see Gross et al., 2013). Filtered data were then resampled to 200 Hz. Electrooculogram artifacts were detected using Independent Component Analysis (ICA) and linearly subtracted from recordings (*fastICA* algorithm implemented in FieldTrip). Artifact rejection was also carried out by excluding all trials with a z-score above the variance threshold of 3. A minimum of 75% artifact-free trials per participant was required for inclusion in subsequent analyses (M = 12, SD = 4.5).

#### Coherence analysis

We used coherence to estimate the phase synchronization between the EEG brain signal and the corresponding auditory stimuli envelopes. Broadband amplitude envelope of the audio signals was obtained from the Hilbert transformed broadband stimulus waveform (Molinaro et al., 2016; Molinaro & Lizarazu, 2018). Then, audio envelopes were resampled from 44100 Hz to 200 Hz. For each experimental condition, coherence between the artifact free epochs and the corresponding audio envelope was calculated in the 0.4 – 15 Hz frequency band with 0.4 Hz frequency resolution (inverse of the trial duration). Following this procedure, we obtained a coherence value for each (i) participant, (ii) condition, (iii) EEG sensor and (iv) frequency bin below 15 Hz.

The coherence bias was estimated empirically for each participant by randomly shuffling the auditory envelopes across trials, and recalculating coherence in 500 permutations with resting state data (Molinaro et al., 2016; der Nederlanden et al., 2022). For each sensor and frequency bin, coherence data were z-score transformed using the mean and standard deviation from the 500 random resting state EEG-audio pairings.

For the subsequent analyses, we extracted the maximum z-scored coherence values of our frequencies of interest from the nearest neighbor bins per each band from each channel for each participant. Figures 3 and 4 bottom show the sensors with the highest coherence values per frequency band and stimuli type. In the Spanish blocks, maximum values were extracted from 1.6 Hz to 2.4 Hz in the case of delta, and 3.6 Hz to 4.4 Hz in the case of theta. In the Basque blocks, maximum values were extracted from 1.2 Hz to 2 Hz in the case of delta, and 3.6 Hz to 4.4 Hz in the case of theta. Trials that deviated by 2 SD or more from the mean were removed from these analyses and are reported in the appendix.

**Figure 3.**
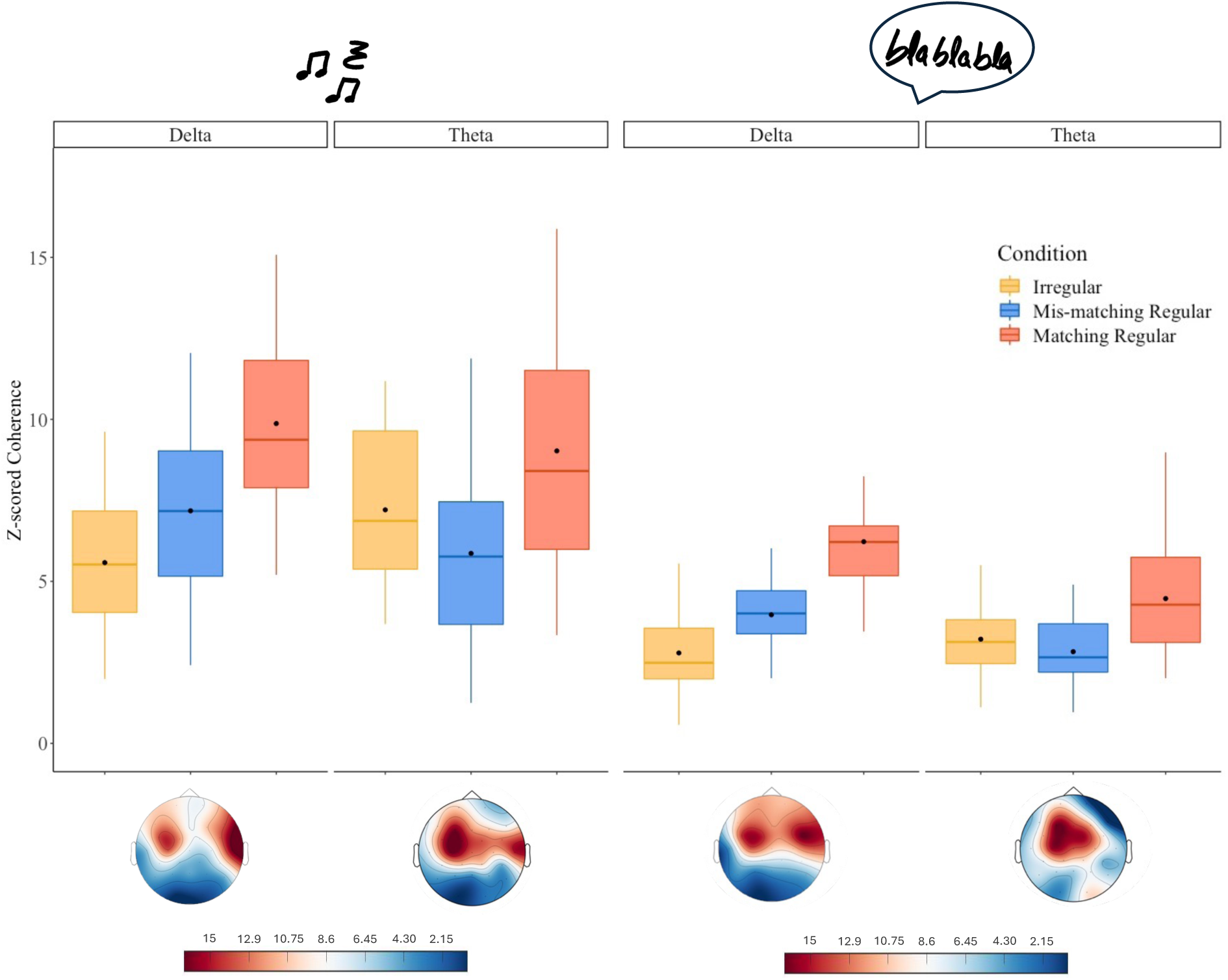
Cortical tracking (z-scores indicating stimulus-brain coherence) of musical and speech sequences in the Spanish blocks. The top left and right panels show the delta and theta frequency bands for music (left) and speech (right). The bottom left and right panels show the topographies of the maximum coherence values extracted for the analysis per frequency and stimulus type.

**Figure 4.**
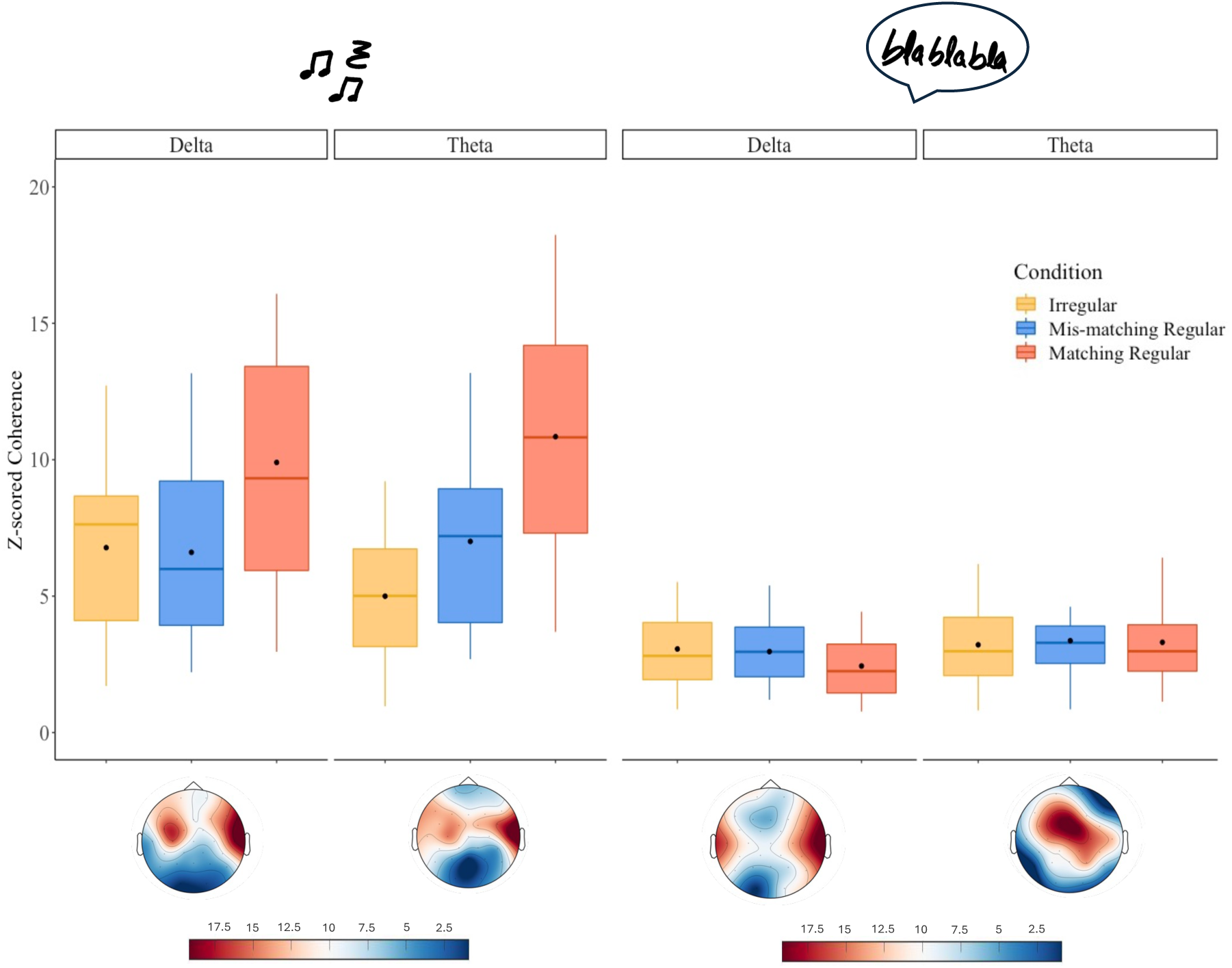
Cortical tracking (z-scores indicating stimulus-brain coherence) of musical and speech sequences in the Basque blocks. The top left and right panels show the delta and theta frequency bands for music (left) and speech (right). The bottom left and right panels show the topographies of the maximum coherence values extracted for the analysis per frequency and stimulus type.

#### Statistical analysis

The main analyses were conducted using Linear mixed effects models constructed with the z-scores transformed maximum coherence values as the dependent variable, including a two-way interaction: Condition (Matching regular, Mis-matching regular, and irregular) x Frequency band (Delta and Theta). As noted above, the Delta and Theta frequency bands were analyzed because they reflect the prosodic and syllabic rhythms of Spanish and Basque. Random intercepts were specified per participant. The continuous variables entered into the analyses were scaled and centered around zero to assist with model convergence. Main effects and interactions were further explored by Post Hoc comparisons using Bonferroni corrections. Analyses were conducted using the lme4 (Bates, 2018) and lmertest (Kuznetsova et al., 2017) packages in R. We also conducted exploratory analyses using the language proficiency and music experience data. These consisted of correlations between z-scored coherence values for each language, stimulus type, frequency bands, condition, participants’ BEST scores, and participants’ individual musical background scores.

## Results

### Behavioral data analysis

Participants’ answers in the behavioral task were scored as correct when the participant correctly identified that the word had or had not appeared in the preceding test block. Accuracy scores were calculated per each condition in each language task. Total number of correct responses was transformed into a percentage. Due to a technical problem with the data collection, we could not collect a complete set of responses from one participant. Overall, participants’ accuracy scores were high (Spanish: M = 87.3 %, SD = 8.9 %; Basque: M = 89.2 %, SD = 8.15 %), indicating that participants were attentive during the task. An ANOVA was run for each language to test if there were any differences in accuracy between the three conditions. No differences between conditions were found in Spanish (*F* (2, 62) = 0.394, *p* = .67) or Basque (*F* (2, 62) = 0.480, *p* = .621).

### EEG analysis

We conducted separate analyses for each language portion of the experiment (Basque and Spanish) and stimulus type (speech sequences and musical sequences). Prior to assessing how exposure to musical sequences with different rhythmic properties impacted cortical tracking of speech, we assessed whether the rhythm manipulation impacted participants’ tracking of music. Therefore, this section is divided into Spanish and Basque, and subdivided into cortical tracking of musical sequences and cortical tracking of speech sequences.

## Spanish

### Cortical tracking of musical sequences

First, we assessed participants’ cortical tracking of musical sequences with different rhythmical properties. Here, we report cortical tracking of musical sequences in the Spanish blocks. The model revealed a main effect of condition (*F* (2, 144) = 25.53, *p* <.001), no main effect of frequency (*F* (1, 144) = 0.186, *p* = .66), and a significant frequency by condition interaction (*F* (2, 144) = 5.486, *p* <.005). The model output can be found in the Appendix, and these results are shown in Figure 3. Post Hoc comparisons revealed that there was higher coherence in the Matching Regular condition than in the Irregular condition (*β* = -.938, *SE* = .147, *t* = −6.392, *p*_bonf_ <.001) and in the Mis-matching Regular condition (*β* = .888, *SE* = .150, *t* = 5.943, *p*_bonf_ <.001). There was no significant difference between the Irregular and Mis-matching regular condition (*β* = -.049, *SE* = .148, *t* = -.333, *p*_bonf_ = 1).

Post Hoc comparisons conducted on the frequency by condition interaction revealed that in the case of delta (1.6 - 2.4 Hz), coherence in the Matching Regular condition was higher than in the Mis-matching (*β* = -.800, *SE* = .216, *t* = 3.699, *p*_bonf_ = .004) and Irregular conditions (*β* = −1.309, *SE* = .212, *t* = −6.182, *p*_bonf_ <.001). There was no difference between the Mis-matching and the Irregular condition (*β* = −.508, *SE* = .213, *t* = −2.384, *p*_bonf_ = .276). In the case of theta (3.6 - 4.4 Hz), coherence in the Matching Regular condition was higher than in the Mis-matching condition (*β* = .976, *SE* = .206, *t* = 4.732, *p*_bonf_ <.001). There was no difference between the Matching Regular condition and the Irregular condition (*β* = −.566, *SE* = .203, *t* = −2.796, *p*_bonf_ = .088), nor the Mis-matching Regular condition and the Irregular condition (*β* = .409, *SE* = .206, *t* = 1.986, *p*_bonf_ = .734).

### Cortical tracking of speech sequences

Next, we assessed cortical tracking of Spanish speech sequences. The model revealed a main effect of condition (*F* (2, 148) = 54.769, *p* <.001), a main effect of frequency (*F* (1, 150) = 16.885, *p* <.001), and a significant frequency by condition interaction (*F* (2, 148) = 11.103 *p* < .001). The model output can be found in the Appendix, and these results are shown in Figure 4. Post Hoc comparisons revealed that there was higher coherence in the Matching Regular condition than in the Irregular condition (*β* = −1.288, *SE* = .133, *t* = - 9.711, *p*_bonf_ <.001) and in the Mis-matching Regular condition (*β* = 1.079, *SE* = .132, *t* = 8.175, *p*_bonf_ <.001). There was no significant difference between the Irregular and Mis-matching Regular condition (*β* = −.209, *SE* = .133, *t* = −.1.573 *p*_bonf_ = .353). Coherence in delta (1.6 – 2.4 Hz) was also significantly higher than coherence in theta (3.6 – 4.4 Hz) (*β* = .445, *SE* = .109, *t* = 4.105, *p*_bonf_ <.001).

### Cortical tracking of speech sequences

Next, we assessed cortical tracking of Spanish speech sequences. The model revealed a main effect of condition (*F* (2, 148) = 54.769, *p* <.001), a main effect of frequency (*F* (1, 150) = 16.885, *p* <.001), and a significant frequency by condition interaction (*F* (2, 148) = 11.103 *p* < .001). The model output can be found in the Appendix, and these results are shown in Figure 3. Post Hoc comparisons revealed that there was higher coherence in the Matching Regular condition than in the Irregular condition (*β* = −1.288, *SE* = .133, *t* = - 9.711, *p*_bonf_ <.001) and in the Mis-matching Regular condition (*β* = 1.079, *SE* = .132, *t* = 8.175, *p*_bonf_ <.001). There was no significant difference between the Irregular and Mis-matching Regular condition (*β* = −.209, *SE* = .133, *t* = −.1.573 *p*_bonf_ = .353). Coherence in delta (1.6 – 2.4 Hz) was also significantly higher than coherence in theta (3.6 – 4.4 Hz) (*β* = .445, *SE* = .109, *t* = 4.105, *p*_bonf_ <.001).

Post Hoc comparisons conducted on the frequency by condition interaction revealed that in the case of delta (1.6 – 2.4 Hz), coherence in the Matching Regular condition was higher than in the Mis-matching Regular (*β* = 1.253, *SE* = .190, *t* = 6.582, *p*_bonf_ <.001) and in the Irregular conditions (*β* = −1.896, *SE* = .189, *t* = −10.021, *p*_bonf_ <.001). Coherence in the Mis-matching Regular condition was significantly higher than in the Irregular condition (*β* = −.642, *SE* = .191, *t* = −3.369, *p*_bonf_ = .014). In the case of theta (3.6 – 4.4 Hz), coherence in the Matching Regular condition was higher than in the Mis-matching Regular (*β* = .903, *SE* = .183, *t* = 4.947, *p*_bonf_ <.001) and in the Irregular conditions (*β* = −.680, *SE* = .186, *t* = −3.658, *p*_bonf_ = .005). There was no difference between the Mis-matching Regular condition and the Irregular condition (*β* = .223, *SE* = .186, *t* = 1.204, *p*_bonf_ = 1)

## Basque

### Cortical tracking of musical sequences

Next, we assessed cortical tracking of musical sequences in the Basque blocks. The model revealed a main effect of condition (*F* (2, 149) = 37.329, *p* <.001), no main effect of frequency (*F* (1, 150) = .158, *p* = .691), and a significant frequency by condition interaction (*F* (2, 149) = 3.180, *p* = .044). The model output can be found in the Appendix, and these results are shown in Figure 4. Post Hoc comparisons revealed that there was higher coherence in the Matching Regular condition than in the Mis-matching Regular condition (*β* = .881, *SE* = .135, *t* = 6.539, *p*_bonf_ <.001) and in the Irregular condition (*β* = −1.104, *SE* = .136, *t* = −8.119, *p*_bonf_ <.001). There was no significant difference between the Irregular and the Mis-matching Regular condition (*β* = −.223, *SE* = .137, *t* = −1.63, *p*_bonf_ = .315).

Post Hoc comparisons conducted on the frequency by condition interaction revealed that in the case of delta (1.2 - 2 Hz), coherence in the Matching Regular condition was higher than in the Mis-matching (*β* = .794, *SE* = .191, *t* = 4.170, *p*_bonf_ <.001) and in the Irregular conditions (*β* = −.772, *SE* = .189, *t* = −4.09, *p*_bonf_ = .001). There was no difference between the Mis-matching Regular condition and the Irregular condition (*β* = .022, *SE* = .193, *t* = .118, *p*_bonf_ = 1). In the case of theta (3.6 – 4.4 Hz), coherence in the Matching Regular condition was higher than in the Mis-matching Regular condition (*β* = .966, *SE* = .190, *t* = 5.078, *p*_bonf_ <.001) and in the Irregular condition (*β* = −1.435, *SE* = .196, *t* = −7.332, *p*_bonf_ <.001). There was no difference between the Mis-matching Regular condition and the Irregular condition (*β* = −.469, *SE* = .194, *t* = 2.417, *p*_bonf_ = .252).

### Cortical tracking of speech sequences

As the next step, we assessed cortical tracking of Basque speech sequences. The model revealed no main effect of condition (*F* (2, 149) = .931, *p* = .396), a main effect of frequency (*F* (1, 152) = 5.952, *p* = .015), and no frequency by condition interaction (*F* (2, 150) = 1.160, *p* = .316). The model output is shown in the Appendix, and these results are shown in Figure 4. Post Hoc comparisons revealed that there was higher coherence in theta (3.6 – 4.4 Hz) than in delta (1.2 - 2 Hz) across conditions (*β* = −.353, *SE* = .145, *t* = −2.437, *p*_bonf_ = .016).

## Interim discussion

In this first experiment, we were interested in clarifying the relevance of the musical-matching rhythms structure on the efficacy of musical stimulation on cortical tracking to speech. In the Spanish blocks, we showed higher cortical tracking of Matching regular sequences compared to Mis-matching and Irregular sequences. In the case of speech, participants showed higher tracking of Spanish speech sequences in the delta band when the sentences were preceded by rhythmical musical cues matched to the temporal structure of the speech (Matching regular sequences). In line with our expectations, participants also showed higher tracking of Mis-matching regular sequences compared to Irregular sequences in the delta band. These results suggest that rhythm regularity was sufficient to enhance cortical tracking of speech in the Spanish blocks.

In the Basque blocks, we found higher cortical tracking of Matching Regular musical sequences in comparison to Mis-matching and Irregular sequences. Participants efficiently tracked the rhythmically different musical sequences that were presented. However, we did not find any influence of the preceding music on the Basque speech sequences, as there was no significant difference between the three conditions. Several explanations were considered for this unexpected finding. First, a lack of the music-to-speech cortical tracking effect could be due to participants’ language knowledge or dominance. However, this is unlikely because participants’ language proficiency scores were high for both languages, and participants were not dominant in one language compared to the other. The possibility of participants not engaging with the Basque task compared to the Spanish task was also considered. However, participants’ behavioral measures showed that participants were attentive to both tasks. Thus, we propose that the inconsistent findings for Spanish and Basque were due to the quality of the language-specific stimuli. The speech stimuli used in this experiment were carefully created controlling for the number of syllables and stressed syllables across languages. Nevertheless, the Basque language allows for some flexibility to position the stress in some words due to the different dialect rules and varieties (Aurrekoetxea et al., 2013). This flexibility allowed us to create stimuli that did not differ that much in the number of syllables and in the rhythmical patterns of the two languages. A later careful inspection of the Basque sentences indicated that in controlling for the position of stressed syllables in the Basque speech sequences, the naturalness of the stimuli was reduced. To test the possibility that this was responsible for the failure to detect a music-to-speech cortical tracking effect in the Basque portion, we created new Basque stimuli allowing for its natural syllable stress position and conducted Experiment 2 including the same participants from Experiment 1.

## Experiment 2

### Method

#### Participants

All 33 participants from Experiment 1 were invited to participate in Experiment 2 and 20 participants were available (6 male, *M* age: 26 years, *SD* = 2.36). In this subsample, all participants were highly proficient in Basque and Spanish (Maximum score: 65; Spanish *M:* 64.12, *SD*: 1.35; Basque *M*: 58.12, *SD* = 6.08), and were actively engaged with music (general musical sophistication factor, maximum score = 100; *M* = 67.89; *SD* = 12.61; 5 participants had more than 5 years of musical formal education). Thus, this subsample was representative of the initial sample from Experiment 1.

The Ethics and Scientific Committee of the BCBL approved the study protocol (approval number 220322ML), which was developed following the declaration of Helsinki. All participants gave their written informed consent prior to the study and received monetary compensation. One participant was excluded due to technical problems during the session, so a total of 19 participants were included in the analysis.

#### Stimuli

The new linguistic stimuli consisted of 64 simple Basque sentences. All were 13 syllables long and were recorded by the same female Basque-Spanish bilingual speaker in natural infant-directed-speech. Each sentence was cued by a metronome to follow a regular rate of 100 bpms, reflecting the rate of stressed syllables in Basque, obtaining a regular meter with an inter-onset interval of 625 msec between accented syllables. Each sentence had a duration of 3750 msec. Identical to Experiment 1, three types of musical stimuli were created: Matching Regular, Mis-matching Regular, and Irregular conditions. The Matching Regular musical sequences reflected the rhythmic structure and pitch contour of linguistic stimuli. The Mis-matching Regular musical sequences preserved the pitch contour of the linguistic stimuli but followed a rhythmical meter with beats that did not match the syllabic structure of the speech stimuli. The Irregular musical sequences were created by altering the pitch and note duration of the Matching Regular sequences and altering their regular meter.

The structure of the task was the same as in Experiment 1. Spectral analyses of the new musical and speech sequences are shown in Figure 5 and revealed peaks at 1.6 Hz and 3.6 Hz.

**Figure 5.**
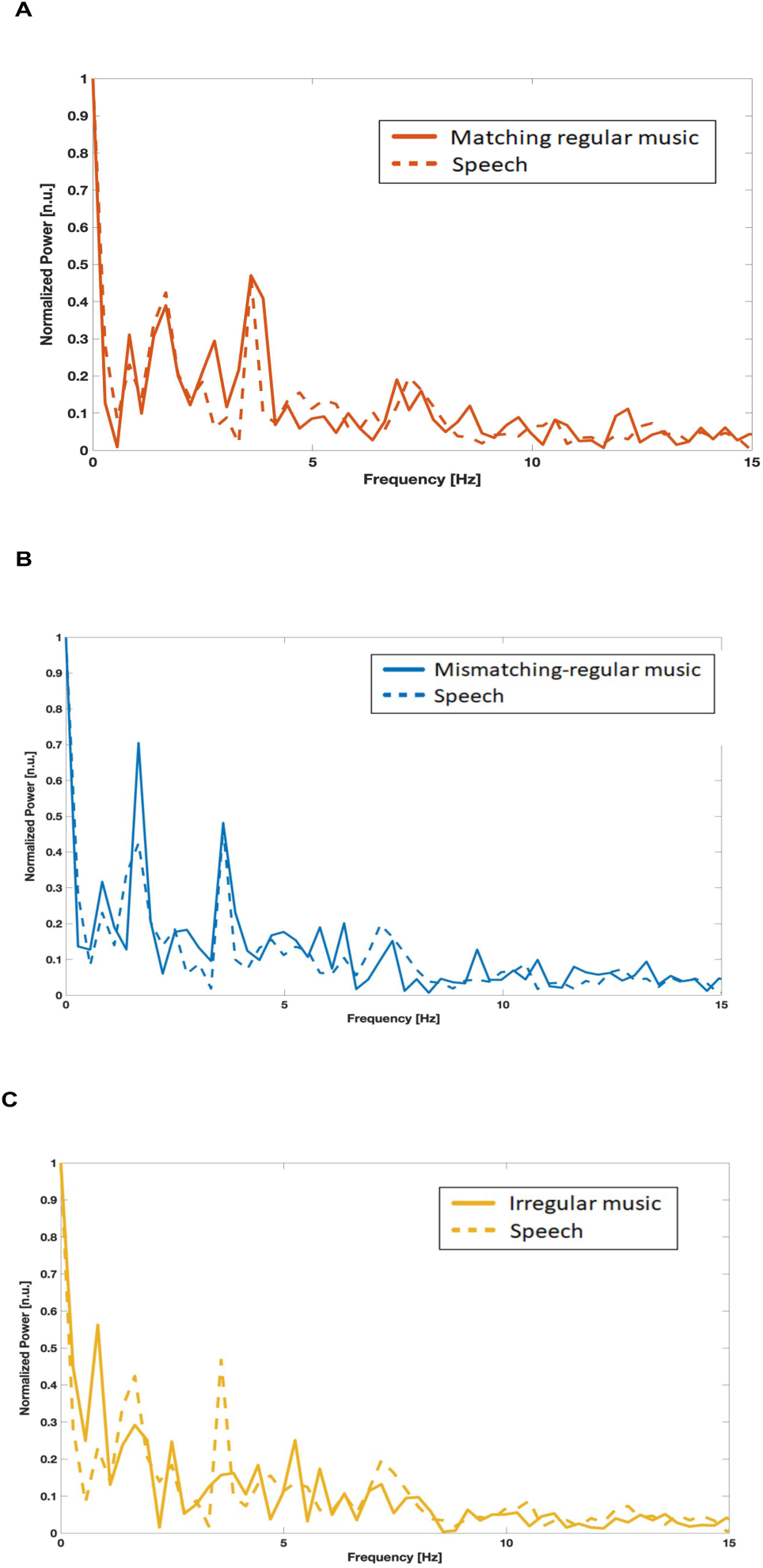
Acoustic properties of Basque musical and speech sequences in Experiment 2. Here we show the spectral information of speech and musical sequences in normalized power (scaled power of signal). Spectra A and B correspond to the new Basque musical stimuli, showing peaks at 1.6 Hz and 3.6 Hz for the Matching and Mis-matching regular condition. Spectrum C corresponds to the new Basque stimuli for the Irregular condition.

#### Procedure

The procedure was identical to Experiment 1 with the exception that participants only completed the Basque portion of the experiment.

#### EEG data acquisition and preprocessing

Data acquisition was the same as in Experiment 1. All processing and analysis steps were identical to the ones from Experiment 1. Given that each sentence was cued by a metronome to follow a regular rate of 100 bpms, the length of the sentences was 3.75 s. Therefore, data were segmented from − 0. s to 3.75 s after the onset of each trial, which gave a total of 384 trials. Since the trial length (3750 msec) led to a frequency resolution of ∼0.26 Hz, we created new trials of 2500 msec with an overlapping window of 1250 msec to achieve a frequency resolution of ∼0.4 Hz for the analyses. The creation of new windows gave a total of 768 trials. A minimum of 75% artifact-free trials per participant was required for inclusion in subsequent analyses (see appendix for trial exclusion summary).

#### Coherence analysis

Coherence was calculated following the steps from Experiment 1. Figure 6 bottom left and right show the sensors with the highest coherence values per frequency band and stimuli type. Maximum values were extracted from 1.2 Hz to 2 Hz in the case of delta, and 3.2 Hz to 4 Hz in the case of theta. Trials that deviated by 2 SD or more from the mean were removed from subsequent analyses (M = 15.3, SD = 3.2).

**Figure 6.**
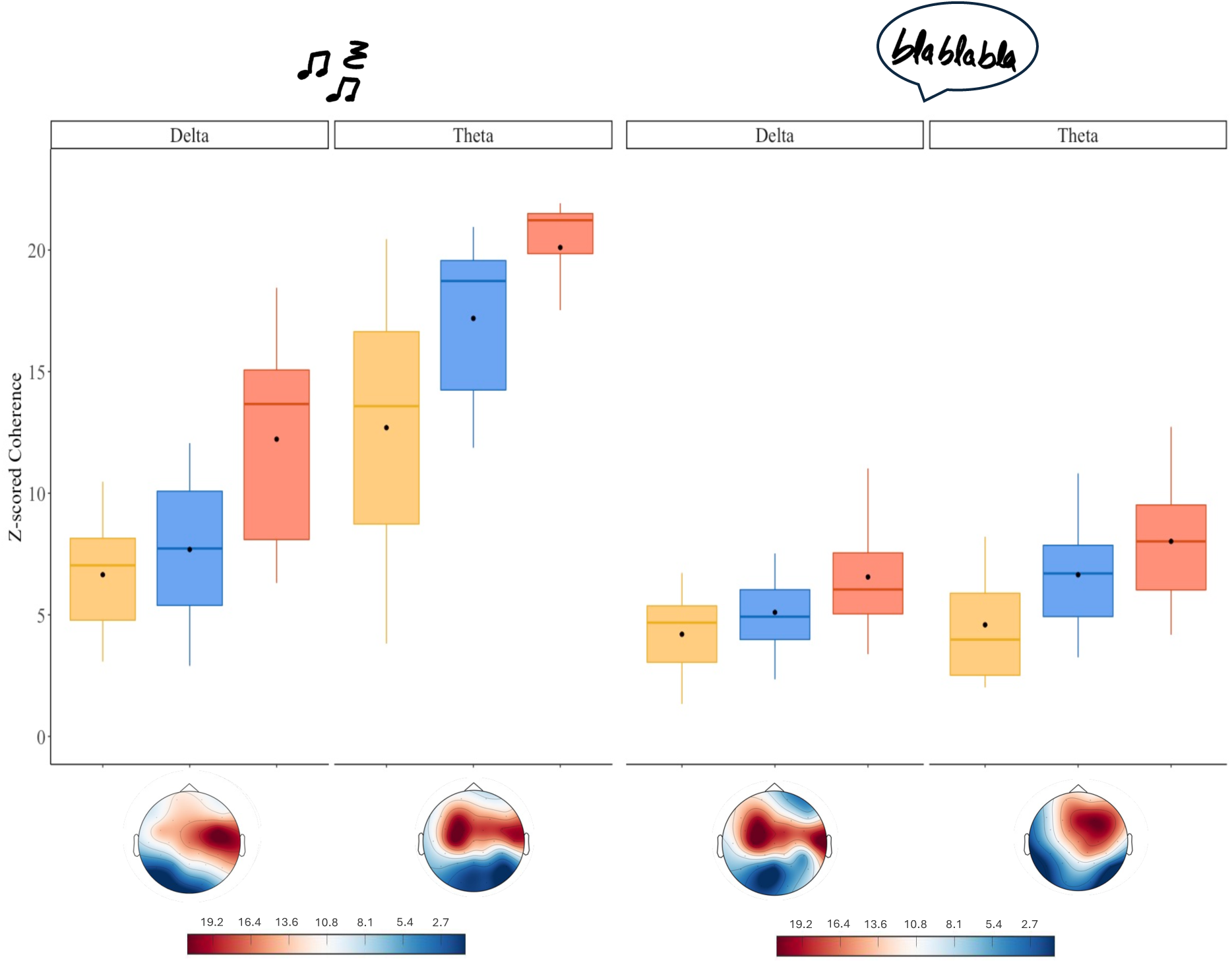
Cortical tracking (z-scores indicating stimulus-brain coherence) of musical and speech sequences in the Basque blocks of Experiment 2. The top left and right panels show the delta and theta frequency bands for music (left) and speech (right). The bottom left and right panels show the topographies of the maximum coherence values extracted for the analysis per frequency and stimulus type.

## Results

### Behavioral data analysis

Participants’ answers in the behavioral task were analyzed as accurate when the participant correctly indicated a word had or had not appeared in the block. Accuracy scores were calculated per condition. Total number of correct responses were transformed into a percentage. Participants’ accuracy scores were high, M = 87.2 %, SD = 6.67 %, indicating that participants were attentive to the task. An ANOVA was run to test differences between conditions. We found a main effect of condition, (*F* (2, 36) = 3.583, *p* = .038). Post-hoc comparisons revealed that participants’ accuracy was higher in the Matching Regular condition compared to the Mis-matching Regular condition, (*p* = .041). There was no difference in accuracy between the Matching Regular condition and the Irregular condition (*p* = .2), nor between the Irregular and the Mismatching Regular condition (*p* = 1).

### EEG analysis

Identical to the first experiment, we conducted separate analyses for each stimulus type (speech and musical sequences). The same analyses were conducted as for Experiment 1.

### Cortical tracking of musical sequences

First, we assessed cortical tracking of musical sequences. The model revealed a main effect of condition (*F* (2, 81) = 53.737, *p* <.001), a main effect of frequency (*F* (1, 82) = 241.352, *p* <.001), and a significant frequency by condition interaction (*F* (2, 81) = 3.3706, *p* = .039). The model output is shown in the Appendix, and these results are shown in Figure 6. Post Hoc comparisons revealed that there was higher coherence in the Matching regular condition than in the Mis-matching Regular condition (*β* = .609, *SE* = .102, *t* = 5.973, *p*_bonf_ <.001) and in the Irregular condition (*β* = −1.037, *SE* = .100, *t* = −10.322, *p*_bonf_ <.001). Coherence was also significantly higher in the Mis-matching regular condition than in the Irregular condition (*β* = −.428, *SE* = .101, *t* = −4.245, *p*_bonf_ <.001). Moreover, there was higher coherence in theta (3.2 - 4 Hz) than in delta (1.2 - 2 Hz) (*β* = −1.28, *SE* = .0827, *t* = −15.528, *p*_bonf_ <.001).

Post Hoc comparisons conducted on the frequency by condition interaction revealed that in the case of delta (1.2 - 2 Hz), there was higher coherence in the Matching Regular condition than in the Mis-matching Regular (*β* = .717, *SE* = .144, *t* = 4.985, *p*_bonf_ <.001) and in the Irregular condition (*β* = −.885, *SE* = .141, *t* = −6.261, *p*_bonf_ <.001). There was no difference in coherence between the Mis-matching Regular and the Irregular condition (*β* = −.168, *SE* = .144, *t* = 1.170, *p*_bonf_ = 1). In the case of theta (3.2 4 Hz), coherence was also significantly higher in the Matching Regular condition than in the Mis-matching Regular (*β* = .501, *SE* = .145, *t* = 3.466, *p*_bonf_ = .012) and in the Irregular condition (*β* = - 1.189, *SE* = .142, *t* = −8.393, *p*_bonf_ <.001). Coherence in the Mis-matching condition was also higher than in the Irregular condition (*β* = −.688, *SE* = .142, *t* = −4.856, *p*_bonf_ <.001).

### Cortical tracking of speech sequences

Next, we assessed cortical tracking of Basque speech sequences. The model revealed a main effect of condition (*F* (2, 83) = 26.268, *p* <.001), a main effect of frequency (*F* (1, 82 = 9.732, *p* = .002), and no significant frequency by condition interaction (*F* (2, 82) = 1.264, *p* = .287). The model output is shown in the Appendix, and these results are shown in Figure 6 Post Hoc comparisons revealed that there was higher coherence in the Matching Regular condition than in the Mis-matching Regular condition (*β* = .551, *SE* = .172, *t* = 3.194, *p*_bonf_ = .005) and in the Irregular condition (*β* = −1.218 *SE* = .169, *t* = - 7.228, *p*_bonf_ <.001). Coherence was also significantly higher in the Mis-matching Regular condition than in the Irregular condition (*β* = −.668, *SE* = .171, *t* = −3.914, *p*_bonf_ <.001). Moreover, there was higher coherence in theta (3.2 - 4 Hz) than in delta (1.2 - 2 Hz) (*β* = −.433, *SE* = .139, *t* = −3.118, *p*_bonf_ = .002).

#### Exploratory analyses: Experiments 1 and 2

We were interested in exploring the relationship between individuals’ musical training, language proficiency, and cortical tracking of speech. For this purpose, we conducted Pearson correlation analyses between the cortical tracking values of musical sequences, speech sequences, participants’ language proficiency scores and participants’ musical background. Given the unexpected findings from the Basque Language in Experiment 1, we decided to include participants’ Spanish scores from Experiment 1 (n = 33) and participants’ Basque scores from Experiment 2 (n = 19). These analyses are reported in Table 8 in the Appendix. Contrary to our expectations, the results of the correlational analyses did not show any consistent relations between musical background and participants’ cortical tracking of music and speech across experiments. Furthermore, there was no evidence of significant relations between the language proficiency scores (BEST scores) in either language or cortical tracking of speech across experiments.

## Discussion

The main aim of this study was to assess whether and how short music exposure might affect subsequent cortical tracking of speech. Specifically, we explored the precise aspects of rhythmic stimulation that lead to an improvement of speech processing through cortical tracking. Secondly, we assessed the effects of language-specific rhythm patterns on cortical tracking of speech in our balanced bilingual sample. To this end, we conducted two experiments where we measured participants’ EEG responses to sentences in Basque and Spanish that were preceded by musical sequences that differed in their beat structure.

In Experiment 1, we found that rhythmical regularities in the Matching and Mis-matching Regular musical sequences were sufficient to enhance cortical tracking of speech sequences in the Spanish language task. Moreover, we showed that when the musical sequences were matched to the meter and beat level, reflecting both the prosodic and syllabic structure of the sentences (Matching Regular), tracking was higher than in the other two conditions. Surprisingly, we did not find this enhancement for the Basque speech sequences. We then conducted Experiment 2 with a subset of the participants, using new stimuli that accurately reflected the prosodic structure of Basque. Consistent with the findings from Experiment 1, we found that rhythmical regularities in the Matching and Mis-matching musical sequences were sufficient to enhance cortical tracking of speech in the Basque task. Again, cortical tracking of speech sequences was higher when the preceding musical sequences shared temporal structure, at the meter and beat level. Overall, these findings suggest that the closer the alignment between the temporal structure of the musical stimuli and the syllabic structure of the speech, the greater the improvement in cortical tracking of the speech sequences.

We will first discuss the findings on cortical tracking of musical sequences and speech sequences. Then, we will discuss the relevance of the musical matching rhythmical structures and, finally, the role of the language-specific rhythms on cortical tracking in relation to our findings.

### Cortical tracking of musical sequences – the speech to music effect

To disentangle the origin of the music-to-speech benefit, it was critical to design musical stimuli that varied in their rhythmicity. To do so, we created musical sequences by converting the prosodic and syllabic structure of the recorded speech to the beat and meter structure of the music (Matching Regular condition). Besides, we then further manipulated the musical sequences to eliminate the beat structure (Mis-Matching Regular condition) and then also the meter structure (Irregular condition). In order to understand whether this difference in rhythmicity could modulate participants’ cortical tracking of the speech sequences, we first inspected participants’ cortical tracking of the musical sequences. Consistent with a wealth of research showing synchronization to rhythmic stimuli (e.g., Fujioka et al., 2009, 2012; Henry and Obleser, 2012; Nozaradan et al., 2011), we showed that participants’ cortical tracking of musical sequences was better when they were rhythmically regular (Matching and Mis-matching Regular) than when they were rhythmically irregular. Surprisingly, despite both the Matching and the Mis-matching musical sequences having a regular meter and structure, participants tracked better the musical sequences that matched the two languages compared to the other two conditions. Based on previous evidence showing that participants effectively track natural music (Harding et al., 2019; der Nederlanden et al., 2020), we expected participants to show effective tracking of both rhythmically regular musical sequences. However, it is important to consider that, although being both highly rhythmical, the main difference between the Matching and Mis-matching conditions was the alignment to the temporal structure of speech. Therefore, these tracking differences may arise from a speech-to-music effect. That is, participants better tracked the musical sequences that best resembled natural speech.

### Cortical tracking to speech sequences – the music to speech effect

One of the explanations for why non-musicians would benefit from short-term musical exposure proposes a shared processing of the structure in language and music (Fiveash et al., 2021; Tierney & Kraus, 2014). The similarities between domains make music an ideal tool to investigate whether this shared structure can be used to enhance speech processing. In this study, we created a shared metrical context between music and speech to investigate the effect of the former on the latter. We provided the first evidence that, by creating musical sequences with a metrical structure based on the prosodic structure of speech, we can use these sequences to modulate how the brain synchronizes to subsequent speech, through the mechanism of cortical tracking. Although a previous study showed that rhythmic scaffolding that matched the syllabic structure of the subsequent speech led to a benefit in cortical tracking of speech (Falk et al., 2017), the specific music properties that best enhance the subsequent synchronization of brain oscillations with speech were unknown. Importantly, we also showed that participants better tracked speech sequences when these were preceded by musical sequences that shared 1) metrical and temporal structure and/or 2) only metrical structure. Our findings suggest that, when presented with a rhythmical stimulus that resembles natural speech, brain oscillations immediately synchronize to the external frequency, supporting the idea that the shared temporal regularity found in music and speech improves synchronization to that regularity.

The present results are significant in three major respects. First, we showed that the regularity found in the rhythmic structure of music acts as a temporal guide for brain oscillations. Previous studies showed that brain oscillations phase-align with the beats of the rhythm (Poeppel & Assaneo, 2020), and that temporal regularity, as found in musical rhythm, could help the brain predict when upcoming beats are coming. Our findings are consistent with the idea that, when presented with rhythmical input, neurons realign the phase of their oscillations so that when an event occurs, they are in their high excitability phase (Schroeder & Lakatos, 2009). This process of alignment has been shown to enhance the neural processing of such events and has sometimes been related to predictive processing. In our case, we provided evidence that a shared temporal framework between speech and music helped maintain the temporal dynamics of the synchronization to the speech signal, even after only very short musical exposure. Our results integrate well with findings supporting the use of short rhythmic exposure, and especially the use of priming paradigms to test improvements of speech perception (Cason & Schön, 2012; Chern et al., 2018; Falk et al., 2017).

Second, our findings suggest that not only the regularity in music is crucial, but so is also adjusting this regularity to optimally reflect the rhythmic characteristics of the language. Although our musical sequences were natural, they were created by extracting the syllabic and melodic patterns of the speech sequences. Therefore, the rhythmical patterns to which we exposed the participants were highly influenced by the speech, that is, they reflected the syllabic and prosodic patterns of each language. In this study, we showed that adjustments to the music play a crucial role on modulating cortical tracking of speech. The importance of closely matching the prosodic characteristics of the music and speech stimuli in order to detect an effect of short music exposure on cortical tracking of speech was corroborated by the need to modify the Basque speech sequences used in Experiment 1 to more accurately reflect the natural prosodic patterns of the language. Importantly, following this adaptation, the results for Basque in Experiment 2 were closely aligned with those for Spanish in Experiment 1.

Third, by testing Basque-Spanish bilinguals in this study, we aimed to identify the role of language-specific rhythms on the music-to-speech cortical tracking effect. Extensive research has shown efficient tracking of listeners’ native languages (see Meyer, 2018 for review), but little is known about the differences in cortical tracking of speech that may arise from cross-linguistic differences at the prosodic and syllabic level. Recent evidence indicates that language-specific features that influence cortical tracking of speech are driven by the specific syllabic and prosodic rates that govern the language rhythms (Peter et al., 2022). While we found a strong effect in both languages, some cross-frequency differences arose for each language.

In Spanish, given its rhythmical characteristics (Molnar et al., 2016; Molnar et al., 2014), we expected to observe a music-to-speech effect mainly in the theta band, based on Falk et al. (2017), who found this effect only in the theta frequency band in speakers of French, a syllabic language. However, while we found similar results when comparing the Matching Regular speech sequences to the two other conditions, we also found an effect in the delta frequency band, where the Matching and the Mismatching regular speech sequences were better tracked than the irregular speech sequences.

It is important to note that our stimuli were recorded in infant-directed speech, and the speaker was cued by a metronome to follow the rate of stressed syllables. The temporal modulation structure of infant-directed speech shows greater modulation energy in the delta frequency band compared to the theta band (Leong et al., 2017). While previous evidence has shown higher cortical tracking to speech in the delta compared to the theta frequency band in Spanish monolingual speakers (Molinaro & Lizarazu, 2018), it is possible that our recordings were mainly influenced by first, the modulations found in infant-directed speech, and second, an emphasis on the prosodic rate.

In the case of Basque, our findings from Experiment 2 showed higher tracking for the Matching and the Mismatching regular speech sequences than the irregular speech sequences in both frequency bands. While few studies have focused on language specific differences in bilinguals’ cortical tracking of speech, Pérez-Navarro et al. (2023) found cortical tracking in both languages only in the delta frequency band but not the theta band on Spanish-Basque bilingual children. In their study, participants’ cortical tracking of speech was modulated by their accumulated experience with the language, being cortical tracking higher for the language they had less exposure with, which was Spanish. Similarly, in a longitudinal study on Spanish-Basque bilingual children, participants showed cortical tracking to Basque speech only in the delta frequency band (Ríos-Lopez et al., 2020). Contrary to Pérez-Navarro et al. (2023), participants’ cortical tracking was not related to participants’ language dominance. In our study, although we did not find any correlation between participants’ language dominance and their cortical tracking measures, it is worth noting that we tested an adult sample that was highly proficient in both Spanish and Basque. It is likely, however, that participants have a greater exposure to one language or the other, something that the BEST language proficiency test does not measure. In another study, Lizarazu et al. (2023) tested Spanish monolinguals that were attending Basque classes. They found that proficiency in Basque modulated participants’ theta-gamma phase-amplitude coupling. This suggests that exposure to the rhythms of one language can influence the cortical tracking of rhythms in another language. In a tapping study, English-French bilinguals’ tapping rates were measured as they tapped to English and French utterances (Lidji et al.,2011). Participants tapped more regularly to English, a stress-timed language, compared to French, a syllable-timed language, suggesting that linguistic experience with a stress-timed language can influence (behavioral) synchronization to speech rhythms. This is an important issue for future research that could inform the developmental trajectory of cortical tracking patterns in multilingual communities.

### Effects of prior musical experience

Finally, we investigated whether musical experience was a modulating factor on the music-to-speech cortical tracking effects we reported here. In a recent study comparing cortical tracking to rhythmically similar spoken utterances and piano melodies, Harding et al. (2019) found greater cortical tracking was significantly related to musical expertise and no intermediate effect was observed for non-musicians. This suggests that musical training, rather than the musical structure itself, predicted cortical tracking of speech. Contrary to expectations, we did not find a significant correlation between cortical tracking of musical sequences and participants’ musical expertise (although, in our study, only 7 participants reported more than 5 years of formal musical education). One potential explanation for this null effect is that the Gold MSI questionnaire we used targets several factors related to musical sophistication, but only one of these factors assesses active musical engagement, including participants’ musical exposure. This factor includes questions about the time spent listening to music and doing other music-related activities, such as going to live concerts. This study was conducted during the COVID 19 pandemic, so scores for the active musical engagement factor might have been lowered as participants were not able to attend music-related activities. Therefore, as a follow-up test, we extracted participants’ musical exposure and correlated just this factor with their cortical tracking of musical sequences. However, there was still no significant correlation. It is maybe even more remarkable that we found a robust effect of short music exposure on cortical tracking of speech, given that this effect was not influenced by participants’ musical background.

#### Limitations and future directions

Our findings raise intriguing questions regarding the effects of acoustic features of music on cortical tracking from a general perspective. In this study we demonstrated that participants’ cortical tracking of speech was enhanced by exposure to rhythmical piano music. However, one issue that was not controlled in this study were differences in the spectral density of the musical sequences (Figures 1 and 7). As can be seen in the spectral density plots, there is a higher peak in the Matching Regular musical sequences for both Spanish and Basque in Experiment 1 in the theta band. However, this limitation does not affect our main finding of more robust music-to-speech effects in the delta frequency band. In addition, although these differences would likely have affected our analyses if we only considered the power of the oscillations, our EEG analyses comprised phase synchronization by the relative amplitude, which are less likely to have been affected.

We still do not know whether participants would also benefit from rhythmical exposure from different acoustic shapes. When considering how oscillations might synchronize to a rhythmic input, it is generally assumed that the behavior should remain consistent across different acoustic forms, whether it’s a sharp pure tone, a percussion beat, or speech. However, little consideration has been paid to how such features might affect the neural ability to synchronize to them (Doelling, 2021). Optimal cortical tracking of different speech rhythms depends on rapid neural responses to large-amplitude acoustic edges or rise times (Huss et al., 2011; Lizarazu et al., 2021), and these edges work as temporal landmarks for cortical tracking to speech. Different instruments have different attack times or rise times, which would lead to differences in their auditory processing. Although how stimulus shape affects neural synchronization and temporal perception is understudied, our results provide further support for the hypothesis that the brain synchronizes to external rhythmic input from piano music. Importantly, while our results show that the rhythmical manipulation is crucial, additional research is needed to better understand how stimulus shape affects the ease with which the brain is able to synchronize to external stimuli, and whether other types of musical stimulation would be better phase-resetters for brain oscillations.

Throughout this paper we have highlighted the importance of the similarities between the speech syllabic and prosodic rhythms and the musical meter and beat structure. However, it is worth noting that together with preserving a regular rhythm, the melodic contour in our study also mimicked the pitch contour of the speech sequences. A limitation of this study is that, given it is one of the main features of prosody, we did not manipulate the pitch contour for the Matching and Mismatching regular musical sequences. However, we did manipulate it for the irregular musical sequences, by scrambling the pitches and the silences of the musical sequences, and thus, distorting the pitch contour based on Falk’s et al. (2017) irregular musical sequences. Future research could address, first, whether preserving the pitch contour but distorting the rhythm of the irregular sequences could influence the music-to-speech cortical tracking effects we found. And second, whether not preserving the pitch contour of speech by adding melody to the musical sequences could result in a null music-to-speech effect.

## Conclusion

Despite the extensive evidence that musical training influences speech processing, the precise aspects that lead to these benefits have not yet been identified. Here, by creating a short-music exposure cross-linguistic paradigm we showed that rhythm, inherent in musical signals, guides the adaptation of brain oscillations. This suggests that cortical tracking is the mechanism through which the brain benefits from music training, by adapting the temporal dynamics of the oscillatory activity to the rhythmic scaffolding of the musical signal. Moreover, we also showed that it is crucial to adjust this rhythm to optimally reflect the rhyhthmic characteristics of the language. These findings are framed in a cross-linguistic context, demonstrating the need to consider the specific rhythms of each language in future studies addressing the music-to-speech cortical tracking effect.

## Data availability

All code used for data processing and analysis available at https://doi.org/10.6084/m9.figshare.22740353.v3. Data available upon request to the authors. All stimuli used for Experiment 1 and 2 are available at https://osf.io/epa4n/?view_only=76177c6fe76b45fbb8eeb33e8446b048

## Acknowledgements

We want to thank the participants for their volunteer contribution to our study. This work was supported by the Basque Government through the BERC 2022-2025 program and Funded by the Spanish State Research Agency through BCBL Severo Ochoa excellence accreditation CEX2020-001010/AEI/10.13039/501100011033. LFM’s work was supported by a Predoctoral Grant from the Spanish Ministry of Science, Innovation and Universities and the European Social Fund, PRE2019-087623. ML’s work was supported by the Ramón y Cajal program of the Spanish MICIU (grant RYC2022-035497-I) and the PIBA_2022_1_0015 from the Basque Government. NM received support from the Spanish Ministry of Science, Innovation and University, (grants PID2022-136991NB-I00; PCI2022-135031-2; PDC2022-133917-I00; RTI2018-096311-B-I0) and from the IKUR initiative. MK’s work was supported by the Spanish State Research Agency through the Ramon y Cajal research fellowship, RYC2018-024284-I.

