## Supplementary material for "Temporal Structure of Music Improves the Cortical Encoding of Speech"

1. Report of trials removal in Experiment 1.

The following number of trials were removed for subsequent analyses since they deviated 2 SD around the mean in each language, per condition and frequency. In the Spanish blocks, regarding the speech sequences in delta: Matching Regular, N = 2, Mis-Matching Regular, N = 1, Irregular, N = 0; in theta: Matching Regular, N = 1, Mis-Matching Regular, N = 2, Irregular, N = 0; musical sequences in delta: Matching Regular, N = 2, Mis-Matching Regular, N = 1, Irregular, N = 2; in theta: Matching Regular, N = 0, Mis-Matching Regular, N = 1, Irregular, N = 0.

2. Report of trials removal in Experiment 2.

The following number of trials were removed for subsequent analyses since they deviated 2 SD around the mean, per condition and frequency: the speech sequences in delta: Matching Regular, N = 1, Mis-Matching Regular, N = 2, Irregular, N = 0; in theta: Matching Regular, N = 2, Mis-Matching Regular, N = 2, Irregular, N = 1; musical sequences in delta: Matching Regular, N = 1, Mis-Matching Regular, N = 2, Irregular, N = 1; in theta: Matching Regular, N = 2, Mis-Matching Regular, N = 2, Irregular, N = 0.

**Table 1**

*Output of the linear mixed effects model on cortical tracking to musical sequences in the Spanish blocks.*

| Effect | ***β*** | *SE* | *df* | *t* | *p* |
| --- | --- | --- | --- | --- | --- |
| Fixed effects |  |  |  |  |  |
| Intercept | -0.570 | .161 | 158 | -3.532 | **<.001** |
| Condition [Matching Regular] | 1.309 | .211 | 144 | 6.186 | **<.001** |
| Condition [Mis-matching Regular] | .508 | .213 | 143 | 2.385 | **.018** |
| Frequency [Theta] | .501 | .206 | 142 | 2.434 | **.016** |
| Condition [Matching Regular] x Frequency [Theta] | −.742 | .029 | 142 | -2.537 | **.012** |
| Condition [Mis-matching Regular] x Frequency [Theta] | -0.918 | .296 | 143 | -3.095 | **.002** |

*Note.* Number of observations = 181, ID (participant) = 33. Model = Music ~ Condition * Frequency + (1|ID)

**Table 2**

*Output of the linear mixed effects model on cortical tracking to Spanish speech sequences.*

| Effect | ***β*** | *SE* | *df* | *t* | *p* |
| --- | --- | --- | --- | --- | --- |
| Fixed effects |  |  |  |  |  |
| Intercept | -.62 | .138 | 173 | -4.471 | **<.001** |
| Condition [Matching Regular] | 1.896 | .189 | 149 | 10.03 | **<.001** |
| Condition [Mis-matching Regular] | .642 | .190 | 149 | 3.371 | **<.001** |
| Frequency [Theta] | .248 | .189 | 149 | 1.315 | .19 |
| Condition [Matching Regular] x Frequency [Theta] | -1.215 | .265 | 149 | -4.587 | **<.001** |
| Condition [Mis-matching Regular] x Frequency [Theta] | -.866 | .266 | 148 | -3.256 | **.001** |

*Note.* Number of observations = 183, ID (participant) = 33. Model = Speech ~ Condition * Frequency + (1|ID)

**Table 3**

*Output of the linear mixed effects model on cortical tracking to musical sequences in the Basque blocks.*

| Effect | ***β*** | *SE* | *df* | *t* | *p* |
| --- | --- | --- | --- | --- | --- |
| Fixed effects |  |  |  |  |  |
| Intercept | -.229 | .156 | 142 | -1.470 | .143 |
| Condition [Matching Regular] | .771 | .188 | 149 | 4.091 | **<.001** |
| Condition [Mis-matching Regular] | -.022 | .192 | 148 | -.118 | .906 |
| Frequency [Theta] | -.429 | .195 | 150 | -2.194 | **.029** |
| Condition [Matching Regular] x Frequency [Theta] | .663 | .271 | 150 | 2.441 | **.015** |
| Condition [Mis-matching Regular] x Frequency [Theta] | .491 | .273 | 149 | 1.798 | .074 |

*Note.* Number of observations = 186, ID (participant) = 33. Model = Music ~ Condition * Frequency + (1|ID)

**Table 4**

*Output of the linear mixed effects model on cortical tracking to Basque speech sequences.*

| Effect | ***β*** | *SE* | *df* | *t* | *p* |
| --- | --- | --- | --- | --- | --- |
| Fixed effects |  |  |  |  |  |
| Intercept | .002 | .180 | 175 | .014 | .989 |
| Condition [Matching Regular] | -.462 | .249 | 150 | -1.852 | .065 |
| Condition [Mis-matching Regular] | -.070 | .251 | 151 | -.279 | .780 |
| Frequency [Theta] | .115 | .249 | 150 | .464 | .643 |
| Condition [Matching Regular] x Frequency [Theta] | .529 | .353 | 149 | 1.499 | .135 |
| Condition [Mis-matching Regular] x Frequency [Theta] | .181 | .354 | 150 | .513 | .608 |

*Note.* Number of observations = 182, ID (participant) = 33. Model = Speech ~ Condition * Frequency + (1|ID)

**Table 5**

*Output of the linear mixed effects model on cortical tracking to musical sequences in the Basque blocks.*

| Effect | ***β*** | *SE* | *df* | *t* | *p* |
| --- | --- | --- | --- | --- | --- |
| Fixed effects |  |  |  |  |  |
| Intercept | -1.008 | .142 | 45 | -7.1 | **<.001** |
| Condition [Matching Regular] | .885 | .141 | 81 | 6.263 | **<.001** |
| Condition [Mis-matching Regular] | .168 | .143 | 81 | 1.17 | .245 |
| Frequency [Theta] | 1.009 | .138 | 81 | 7.263 | **<.001** |
| Condition [Matching Regular] x Frequency [Theta] | .303 | .199 | 81 | 1.524 | .131 |
| Condition [Mis-matching Regular] x Frequency [Theta] | .519 | .201 | 81 | 2.577 | .011 |

*Note.* Number of observations = 106, ID (participant) = 19. Model = Music ~ Condition * Frequency + (1|ID)

**Table 6**

*Output of the linear mixed effects model on cortical tracking to Basque speech sequences.*

| Effect | ***β*** | *SE* | *df* | *t* | *p* |
| --- | --- | --- | --- | --- | --- |
| Fixed effects |  |  |  |  |  |
| Intercept | -.661 | .196 | 68 | -3.360 | **.001** |
| Condition [Matching Regular] | .992 | .234 | 82 | 4.231 | **<.001** |
| Condition [Mis-matching Regular] | .431 | .238 | 82 | 1.809 | .07 |
| Frequency [Theta] | .125 | .234 | 82 | .534 | .595 |
| Condition [Matching Regular] x Frequency [Theta] | .452 | .337 | 82 | 1.342 | .183 |
| Condition [Mis-matching Regular] x Frequency [Theta] | .472 | .339 | 82 | 1.393 | .167 |

*Note.* Number of observations = 106, ID (participant) = 19. Model = Speech ~ Condition * Frequency + (1|ID)

**Table 7. Pearson Correlation analyses between between the cortical tracking values of musical sequences, speech sequences, participants’ language proficiency scores and participants’ musical background.**

**Experiment 1.**

| **Pearson's Correlations** | | | | | | | | | | | | | | | | | | | | | | | | | | | | | | | | | | | | | | | | |
| --- | --- | --- | --- | --- | --- | --- | --- | --- | --- | --- | --- | --- | --- | --- | --- | --- | --- | --- | --- | --- | --- | --- | --- | --- | --- | --- | --- | --- | --- | --- | --- | --- | --- | --- | --- | --- | --- | --- | --- | --- |
| **Variable** | |  | | **BEST MARK 65 SPANISH** | | **Active Musical Engagement** | | **Perceptual Abilities** | | **Musical Training** | | **Singing Abilities** | | **Emotional Engagement with music** | | **General Musical Sophistication** | | **Tracking to speech - MR - delta** | | **Tracking to speech - MM - delta** | | **Tracking to speech - IR - delta** | | **Tracking to speech - MR - theta** | | **Tracking to speech - MM - theta** | | **Tracking to speech - IR - theta** | | **Tracking to music - MR - delta** | | **Tracking to music - MM - delta** | | **Tracking to music - IR - delta** | | **Tracking to music - MR - theta** | | **Tracking to music - MM - theta** | | **Tracking to music - IR - theta** |
| 1. BEST MARK 65 SPANISH |  |  |  | — |  |  |  |  |  |  |  |  |  |  |  |  |  |  |  |  |  |  |  |  |  |  |  |  |  |  |  |  |  |  |  |  |  |  |  |  |
| 2. Active Musical Engagement |  |  |  | -0.101 |  | — |  |  |  |  |  |  |  |  |  |  |  |  |  |  |  |  |  |  |  |  |  |  |  |  |  |  |  |  |  |  |  |  |  |  |
| 3. Perceptual Abilities |  |  |  | -0.012 |  | 0.249 |  | — |  |  |  |  |  |  |  |  |  |  |  |  |  |  |  |  |  |  |  |  |  |  |  |  |  |  |  |  |  |  |  |  |
| 4. Musical Training |  |  |  | -0.010 |  | 0.118 |  | 0.265 |  | — |  |  |  |  |  |  |  |  |  |  |  |  |  |  |  |  |  |  |  |  |  |  |  |  |  |  |  |  |  |  |
| 5. Singing Abilities |  |  |  | -0.173 |  | 0.561*** |  | 0.377* |  | -0.028 |  | — |  |  |  |  |  |  |  |  |  |  |  |  |  |  |  |  |  |  |  |  |  |  |  |  |  |  |  |  |
| 6. Emotional Engagement with music |  |  |  | -0.120 |  | 0.269 |  | 0.699*** |  | 0.361* |  | 0.152 |  | — |  |  |  |  |  |  |  |  |  |  |  |  |  |  |  |  |  |  |  |  |  |  |  |  |  |  |
| 7. General Musical Sophistication |  |  |  | -0.085 |  | 0.547*** |  | 0.738*** |  | 0.595*** |  | 0.332 |  | 0.843*** |  | — |  |  |  |  |  |  |  |  |  |  |  |  |  |  |  |  |  |  |  |  |  |  |  |  |
| 8. Tracking to speech - MR - delta |  |  |  | 0.240 |  | -0.008 |  | -0.067 |  | -0.068 |  | 0.041 |  | 0.017 |  | -0.021 |  | — |  |  |  |  |  |  |  |  |  |  |  |  |  |  |  |  |  |  |  |  |  |  |
| 9. Tracking to speech - MM - delta |  |  |  | 0.065 |  | -0.055 |  | -0.028 |  | 0.023 |  | 0.143 |  | -0.158 |  | -0.067 |  | 0.439* |  | — |  |  |  |  |  |  |  |  |  |  |  |  |  |  |  |  |  |  |  |  |
| 10. Tracking to speech - IR - delta |  |  |  | -0.140 |  | 0.014 |  | 0.037 |  | 0.023 |  | -0.016 |  | 0.130 |  | 0.090 |  | 0.259 |  | 0.346 |  | — |  |  |  |  |  |  |  |  |  |  |  |  |  |  |  |  |  |  |
| 11. Tracking to speech - MR - theta |  |  |  | -0.098 |  | 0.224 |  | 0.092 |  | -0.301 |  | 0.299 |  | -0.012 |  | 0.019 |  | 0.169 |  | -0.058 |  | 0.088 |  | — |  |  |  |  |  |  |  |  |  |  |  |  |  |  |  |  |
| 12. Tracking to speech - MM - theta |  |  |  | -0.070 |  | 0.374* |  | 0.026 |  | -0.167 |  | 0.005 |  | 0.120 |  | 0.195 |  | -0.237 |  | -0.177 |  | 0.054 |  | 0.356* |  | — |  |  |  |  |  |  |  |  |  |  |  |  |  |  |
| 13. Tracking to speech - IR - theta |  |  |  | 0.032 |  | 0.171 |  | 0.331 |  | 0.045 |  | 0.215 |  | 0.264 |  | 0.252 |  | -0.042 |  | -0.183 |  | -0.343 |  | 0.362 |  | -0.003 |  | — |  |  |  |  |  |  |  |  |  |  |  |  |
| 14. Tracking to music - MR - delta |  |  |  | 0.074 |  | 0.279 |  | 0.023 |  | 0.224 |  | 0.100 |  | 0.242 |  | 0.305 |  | 0.147 |  | -0.076 |  | 0.109 |  | -0.170 |  | 0.185 |  | -0.255 |  | — |  |  |  |  |  |  |  |  |  |  |
| 15. Tracking to music - MM - delta |  |  |  | -0.090 |  | 0.176 |  | 0.253 |  | -0.095 |  | 0.410 | * | 0.173 |  | 0.171 |  | -0.207 |  | -0.046 |  | 0.183 |  | 0.102 |  | 0.378 |  | -0.254 |  | 0.298 |  | — |  |  |  |  |  |  |  |  |
| 16. Tracking to music - IR - delta |  |  |  | 0.008 |  | 0.089 |  | -0.154 |  | -0.169 |  | -0.034 |  | -0.137 |  | -0.140 |  | -0.213 |  | -0.273 |  | -0.022 |  | 0.223 |  | 0.283 |  | -0.013 |  | -0.021 |  | 0.188 |  | — |  |  |  |  |  |  |
| 17. Tracking to music - MR - theta |  |  |  | 0.297 |  | 0.204 |  | -0.050 |  | -0.001 |  | 0.054 |  | -0.031 |  | 0.021 |  | 0.463* |  | 0.416* |  | 0.259 |  | 0.165 |  | 0.004 |  | 0.064 |  | -0.143 |  | 0.024 |  | 0.307 |  | — |  |  |  |  |
| 18. Tracking to music - MM - theta |  |  |  | -0.157 |  | 0.335 |  | -0.241 |  | -0.063 |  | -0.131 |  | 0.029 |  | 0.033 |  | 0.204 |  | 0.035 |  | 0.355 |  | 0.213 |  | 0.341 |  | -0.073 |  | 0.189 |  | 0.118 |  | 0.458* |  | 0.539** |  | — |  |  |
| 19. Tracking to music - IR - theta |  |  |  | 0.202 |  | -0.091 |  | -0.221 |  | -0.062 |  | -0.091 |  | -0.077 |  | -0.161 |  | 0.483** |  | 0.225 |  | 0.013 |  | 0.009 |  | -0.173 |  | -0.061 |  | -0.083 |  | -0.167 |  | 0.084 |  | 0.325 |  | 0.186 |  | — |
| *** p < .05, ** p < .01, *** p < .001** | | | | | | | | | | | | | | | | | | | | | | | | | | | | | | | | | | | | | | | | |

**Experiment 2.**

| **Pearson's Correlations** | | | | | | | | | | | | | | | | | | | | | | | | | | | | | | | | | | | | | | | | |
| --- | --- | --- | --- | --- | --- | --- | --- | --- | --- | --- | --- | --- | --- | --- | --- | --- | --- | --- | --- | --- | --- | --- | --- | --- | --- | --- | --- | --- | --- | --- | --- | --- | --- | --- | --- | --- | --- | --- | --- | --- |
| **Variable** | |  | | **BEST MARK 65 BASQUE** | | **Active Musical Engagement** | | **Perceptual Abilities** | | **Musical Training** | | **Singing Abilities** | | **Emotional Engagement with music** | | **General Musical Sophistication** | | **Tracking to speech - MR - delta** | | **Tracking to speech - MM - delta** | | **Tracking to speech - IR - delta** | | **Tracking to speech - MR - theta** | | **Tracking to speech - MM - theta** | | **Tracking to speech - IR - theta** | | **Tracking to music - MR - delta** | | **Tracking to music - MM - delta** | | **Tracking to music - IR - delta** | | **Tracking to music - MR - theta** | | **Tracking to music - MM - theta** | | **Tracking to music - IR - theta** |
| 1. BEST MARK 65 BASQUE |  |  |  | — |  |  |  |  |  |  |  |  |  |  |  |  |  |  |  |  |  |  |  |  |  |  |  |  |  |  |  |  |  |  |  |  |  |  |  |  |
| 2. Active Musical Engagement |  |  |  | 0.188 |  | — |  |  |  |  |  |  |  |  |  |  |  |  |  |  |  |  |  |  |  |  |  |  |  |  |  |  |  |  |  |  |  |  |  |  |
| 3. Perceptual Abilities |  |  |  | -0.078 |  | 0.323 |  | — |  |  |  |  |  |  |  |  |  |  |  |  |  |  |  |  |  |  |  |  |  |  |  |  |  |  |  |  |  |  |  |  |
| 4. Musical Training |  |  |  | 0.144 |  | 0.335 |  | 0.214 |  | — |  |  |  |  |  |  |  |  |  |  |  |  |  |  |  |  |  |  |  |  |  |  |  |  |  |  |  |  |  |  |
| 5. Singing Abilities |  |  |  | 0.219 |  | 0.586** |  | 0.436 |  | -0.112 |  | — |  |  |  |  |  |  |  |  |  |  |  |  |  |  |  |  |  |  |  |  |  |  |  |  |  |  |  |  |
| 6. Emotional Engagement with music |  |  |  | -0.188 |  | 0.310 |  | 0.757*** |  | 0.450 |  | 0.090 |  | — |  |  |  |  |  |  |  |  |  |  |  |  |  |  |  |  |  |  |  |  |  |  |  |  |  |  |
| 7. General Musical Sophistication |  |  |  | -0.015 |  | 0.623** |  | 0.753*** |  | 0.674** |  | 0.338 |  | 0.856*** |  | — |  |  |  |  |  |  |  |  |  |  |  |  |  |  |  |  |  |  |  |  |  |  |  |  |
| 8. Tracking to speech - MR - delta |  |  |  | 0.183 |  | 0.212 |  | -0.420 |  | -0.230 |  | -0.141 |  | -0.331 |  | -0.297 |  | — |  |  |  |  |  |  |  |  |  |  |  |  |  |  |  |  |  |  |  |  |  |  |
| 9. Tracking to speech - MM - delta |  |  |  | -0.114 |  | 0.043 |  | 0.015 |  | -0.337 |  | -0.217 |  | 0.146 |  | -0.109 |  | 0.512* |  | — |  |  |  |  |  |  |  |  |  |  |  |  |  |  |  |  |  |  |  |  |
| 10. Tracking to speech - IR - delta |  |  |  | -0.095 |  | 0.437 |  | 0.049 |  | 0.216 |  | 0.141 |  | 0.235 |  | 0.323 |  | 0.238 |  | 0.164 |  | — |  |  |  |  |  |  |  |  |  |  |  |  |  |  |  |  |  |  |
| 11. Tracking to speech - MR - theta |  |  |  | 0.439 |  | 0.304 |  | 0.352 |  | 0.082 |  | 0.374 |  | 0.128 |  | 0.343 |  | 0.134 |  | -0.197 |  | 0.063 |  | — |  |  |  |  |  |  |  |  |  |  |  |  |  |  |  |  |
| 12. Tracking to speech - MM - theta |  |  |  | 0.042 |  | 0.394 |  | 0.216 |  | 0.153 |  | 0.317 |  | 0.227 |  | 0.416 |  | 0.284 |  | -0.111 |  | 0.213 |  | 0.763*** |  | — |  |  |  |  |  |  |  |  |  |  |  |  |  |  |
| 13. Tracking to speech - IR - theta |  |  |  | 0.452 |  | 0.330 |  | 0.207 |  | 0.118 |  | 0.366 |  | 0.075 |  | 0.222 |  | 0.377 |  | -0.104 |  | 0.227 |  | 0.807*** |  | 0.673** |  | — |  |  |  |  |  |  |  |  |  |  |  |  |
| 14. Tracking to music - MR - delta |  |  |  | 0.380 |  | 0.019 |  | -0.059 |  | 0.042 |  | -0.008 |  | -0.020 |  | 0.016 |  | 0.408 |  | 0.254 |  | -0.080 |  | 0.255 |  | 0.141 |  | 0.326 |  | — |  |  |  |  |  |  |  |  |  |  |
| 15. Tracking to music - MM - delta |  |  |  | 0.248 |  | 0.007 |  | 0.081 |  | -0.165 |  | -0.084 |  | 0.036 |  | 8.069×10^-4^ |  | 0.371 |  | 0.016 |  | -0.122 |  | 0.506 |  | 0.270 |  | 0.234 |  | 0.552* |  | — |  |  |  |  |  |  |  |  |
| 16. Tracking to music - IR - delta |  |  |  | 0.170 |  | -0.004 |  | -0.022 |  | -0.123 |  | -0.055 |  | -0.170 |  | -0.118 |  | 0.286 |  | 0.092 |  | 0.082 |  | 0.503* |  | 0.107 |  | 0.324 |  | 0.668** |  | 0.858*** |  | — |  |  |  |  |  |  |
| 17. Tracking to music - MR - theta |  |  |  | 0.317 |  | 0.181 |  | 0.338 |  | -0.084 |  | 0.185 |  | 0.186 |  | 0.213 |  | 0.221 |  | 0.389 |  | 0.025 |  | 0.531* |  | 0.256 |  | 0.369 |  | 0.342 |  | 0.393 |  | 0.254 |  | — |  |  |  |  |
| 18. Tracking to music - MM - theta |  |  |  | 0.306 |  | -0.080 |  | 0.144 |  | 0.011 |  | -0.158 |  | -0.041 |  | 3.367×10^-4^ |  | 0.120 |  | 0.163 |  | -0.425 |  | 0.526* |  | 0.181 |  | 0.432 |  | 0.327 |  | 0.415 |  | 0.310 |  | 0.616** |  | — |  |  |
| 19. Tracking to music - IR - theta |  |  |  | 0.165 |  | -0.126 |  | 0.030 |  | -0.119 |  | -0.237 |  | -0.031 |  | -0.104 |  | 0.231 |  | 0.264 |  | -0.322 |  | 0.411 |  | 0.057 |  | 0.114 |  | 0.431 |  | 0.751*** |  | 0.655** |  | 0.480 |  | 0.407 |  | — |
| * p < .05, ** p < .01, *** p < .001 | | | | | | | | | | | | | | | | | | | | | | | | | | | | | | | | | | | | | | | | |
